# A β-catenin-driven switch in TCF/LEF transcription factor binding to DNA target sites promotes commitment of mammalian nephron progenitor cells

**DOI:** 10.1101/2020.10.26.356410

**Authors:** Qiuyu Guo, Albert Kim, Bin Li, Andrew Ransick, Helena Bugacov, Xi Chen, Nils Lindstrom, Aaron Brown, Leif Oxburgh, Bing Ren, Andrew P. McMahon

**Affiliations:** Department of Stem Cell Biology and Regenerative Medicine, Eli and Edythe Broad-CIRM Center for Regenerative Medicine and Stem Cell Research, Keck School of Medicine of the University of Southern California, CA 90089, USA; Center for Molecular Medicine, Maine Medical Center Research Institute, 81 Research Drive, Scarborough, ME 04074, USA; The Rogosin Institute, New York, NY, 10065, USA; Ludwig Institute for Cancer Research, Department of Cellular and Molecular Medicine, Institute of Genomic Medicine, Moores Cancer Center, University of California San Diego, La Jolla, California, United States of America

## Abstract

The canonical Wnt pathway transcriptional co-activator β-catenin regulates self-renewal and differentiation of mammalian nephron progenitor cells (NPCs). We modulated β-catenin levels in NPC cultures using the GSK3 inhibitor CHIR9902 (CHIR) to examine opposing developmental actions of β-catenin. Low CHIR-mediated maintenance and expansion of NPCs is independent of direct engagement of TCF/LEF/ β-catenin transcriptional complexes at low-CHIR dependent cell-cycle targets. In contrast, in high CHIR, TCF7/LEF1/β-catenin complexes replaced TCF7L1/TCF7L2 binding on enhancers of differentiation-promoting target genes. Chromosome confirmation studies showed pre-established promoter-enhancer connections to these target genes in NPCs. High CHIR-associated *de novo* looping was observed in positive transcriptional feedback regulation to the canonical Wnt pathway. Thus, β-catenin’s direct transcriptional role is restricted to the induction of NPCs where rising β-catenin levels switch inhibitory TCF7L1/TCF7L2 complexes to activating LEF1/TCF7 complexes at primed gene targets poised for rapid initiation of a nephrogenic program.

## Introduction

The now classical model of canonical Wnt signaling invokes two transcriptional states (Wiese, Nusse, and van Amerongen 2018, Steinhart and Angers 2018). In the absence of Wnt ligand, HMG box family Tcf transcription factors bind enhancers of Wnt target genes recruiting co-repressors (Tle, Ctbp and others) to silence target gene expression. In the cytoplasm, the transcriptional co-activator β-catenin is phosphorylated by an axin/GSK3ß-dependent β-catenin destruction complex, resulting in ubiquitin-mediated proteasomal degradation. Upon Wnt ligand binding to Fzd receptor/Lrp co-receptors on the cell surface, the β-catenin destruction complex is sequestered to the activated receptor protein complex through axin interactions, removing β-catenin from GSK3ß-directed, phosphorylation-mediated degradation (Schaefer and Peifer 2019). As a result of increasing β-catenin levels, β-catenin is free to associate with TCF/LEF DNA binding partners, activating Wnt target gene transcription (Mosimann, Hausmann, and Basler 2009). While evidence suggests all four mammalian Tcf family members are able to functionally interact with both Tle family co-repressors and β-catenin (Brantjes et al. 2001), a variety of studies in a range of biological systems indicate that Tcf7l1 predominantly acts as a repressor, Tcf7l2 as a context-dependent activator or repressor, and Tcf7 and Lef1 as transcriptional activators, of Wnt target gene expression (Lien and Fuchs 2014).

The adult (metanephric) mammalian kidney arises from distinct cell populations within the intermediate mesoderm (McMahon 2016). All nephrons – the repeating functional unit of the kidney – arise from a small pool of a few hundred nephron progenitor cells (NPCs) established at the onset of kidney development (Short et al. 2014, Kobayashi et al. 2008). The subsequent balance in the maintenance, expansion and commitment of NPCs is critical to ensuring a full complement of nephrons, approximately 14,000 in the mouse and 1 million in the human kidney (Bertram et al. 2011). A reduced nephron endowment has been associated with abnormal kidney function and disease susceptibility (Luyckx and Brenner 2010, McMahon 2016, Bertram et al. 2011). The maintenance and expansion of NPCs is supported by Fgf, Bmp and Wnt signals produced by NPCs or adjacent mesenchymal interstitial progenitor and ureteric epithelial cell types (McMahon 2016). Within this nephrogenic niche, ureteric epithelium-derived Wnt9b is thought to act on NPCs in a β-catenin-dependent transcriptional process to regulate NPC target gene expression and expansion of the nephron progenitor pool (Karner et al. 2011). The removal of *Wnt9b* from the ureteric epithelium and NPC-specific production of β-catenin also results in the failure of NPC differentiation (Carroll et al. 2005), whereas chemical inhibition of GSK3β (Davies and Garrod 1995, Kuure et al. 2007), or genetic activation within NPCs of a β-catenin form insensitive to GSK phosphorylation-mediated proteasomal degradation, leads to Wnt9b-independent ectopic induction of differentiation-promoting gene targets (Park, Valerius, and McMahon 2007). Genomic analysis of β-catenin engagement at TCF/LEF recognition motifs within enhancers linked to genes driving NPC differentiation (Park et al. 2012), and subsequent transgenic studies demonstrating TCF/LEF-dependent activity of cis regulatory elements, provide strong evidence for a canonical Wnt/β-catenin/Tcf regulatory axis (Mosimann, Hausmann, and Basler 2009). Thus, canonical Wnt signaling directs opposing NPC programs: maintenance and expansion of uncommitted NPCs and their commitment to nephron formation.

In this study, we employed an *in vitro* model to investigate the genomic regulatory mechanisms underlying the diverse action of canonical Wnt signaling in NPC programs. In this system, maintenance and expansion of NPCs, or their commitment to a nephrogenic program, are controlled by varying levels of CHIR99021 (Cohen and Goedert, 2004; abbreviated to ‘CHIR’ in the following text) addition to a chemically-defined nephron progenitor expansion medium (NPEM) (Brown, Muthukrishnan, and Oxburgh 2015). CHIR binding to Gsk3β inhibits Gsk3β-mediated phosphorylation and proteasomal degradation of β-catenin (Yost et al. 1996, Aberle et al. 1997). Analysis of chromatin interactions and TCF/LEF factor engagement at DNA targets support a model where β-catenin levels act as a key regulatory switch to modify TCF/LEF complex engagement at DNA targets and commitment of NPCs to a nephron forming program.

## Results

### Elevated CHIR levels mediate a rapid inductive response in mouse nephron progenitor cells

A low level of CHIR (1.25 μM) is an essential component in NPEM medium supporting the expansion of NPCs while maintaining the nephron-forming competence (Brown et al., 2015). Within 3 days of elevating CHIR levels (3 μM), aggregate NPC cultures show a robust signature of nephron differentiation (Brown, Muthukrishnan, and Oxburgh 2015). To develop this system further for detailed molecular characterization of CHIR/β-catenin-directed transcriptional events, we collected NPCs from E16.5 embryonic kidneys by magnetic-activated cell sorting (MACS; (Brown, Muthukrishnan, and Oxburgh 2015). NPCs were cultured in NPEM supplemented with a maintenance level of CHIR (1.25 μM – low CHIR thoughout) to promote self-renewal of NPCs. CHIR levels where then titrated to determine an effective concentration for a rapid activation of early target genes of NPC commitment, mirroring *in vivo* responses.

As expected, low CHIR conditions maintained Six2, a key determinant of the NPC state, but did not induce expression of Jag1 (Fig 1B-C, Fig S1B), a Notch pathway ligand activity at early stage of mouse and human NPC commitment (Georgas et al. 2009, Lindstrom, McMahon, et al. 2018). A significant increase of cellular and nuclear β-catenin (Fig 1A, B and Fig S3C) was observed in 5 μM CHIR (‘high CHIR’ throughout), along with a strong inductive response: Six2 protein level persisted, but there was a robust induction of Jag1(Fig 1A-B), mirroring early inductive events in the pre-tubular aggregate and renal vesicle *in vivo* (Lindstrom, McMahon, et al. 2018, Mugford et al. 2009, Georgas et al. 2009, Xu, Liu, et al. 2014). We adopted this induction condition throughout the study.

**Figure 1.**
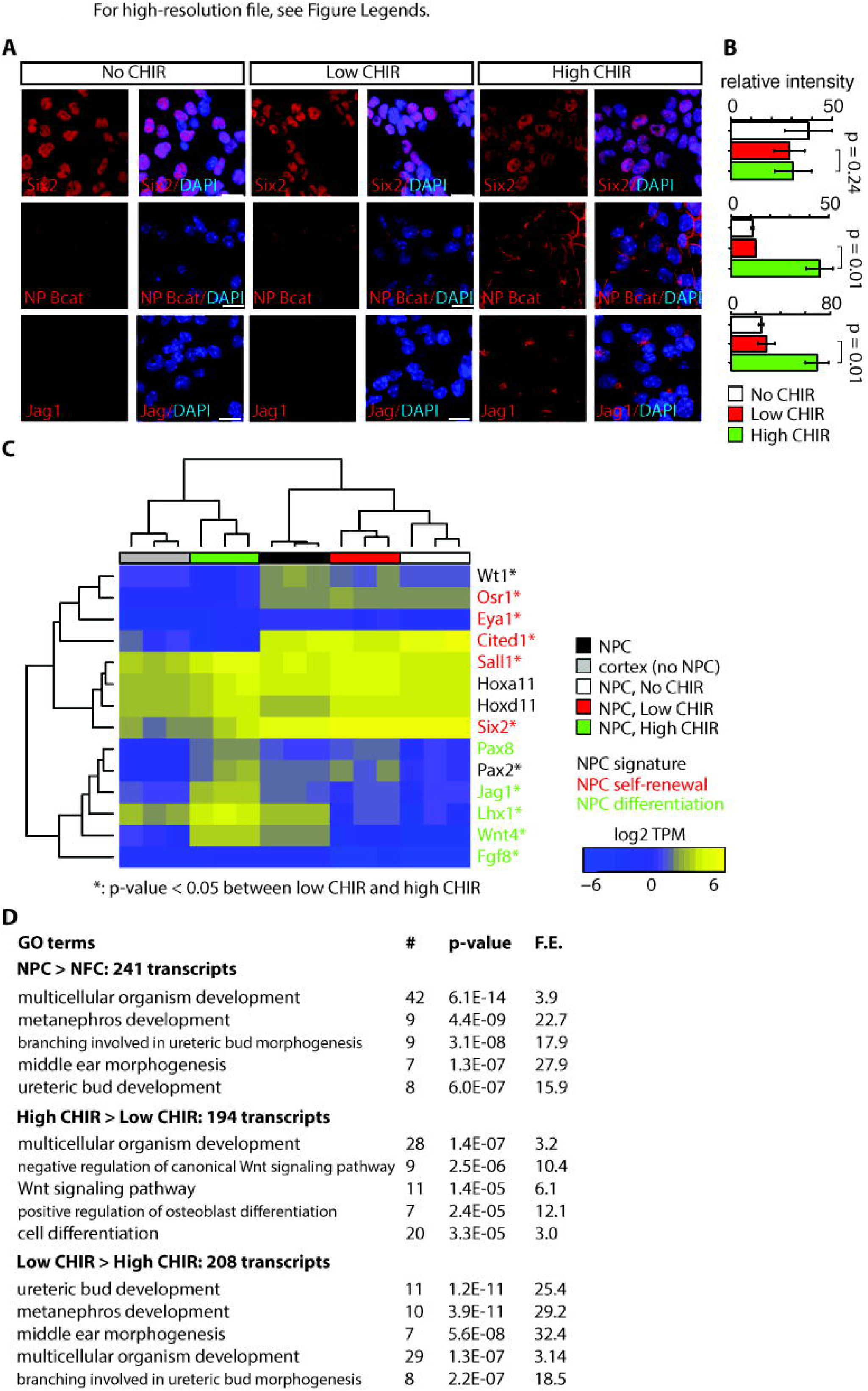
NPEM supplemented with differential levels of CHIR99021 models nephron progenitor cell maintenance or differentiation in a plate. (A) Immuno-fluorescence (IF) staining showing expression level of Six2, non-phospho (NP) β-catenin and Jag1 in NPC cultured in NPEM supplemented with various CHIR dosages. (B) Relative intensity of IF signals from individual cells in experiment associated with A. (C) Heatmap/Hierachical cluster of expression levels of NPC signature, self-renewal and differentiation marker genes. (D) Top 5 enriched GO terms of indicated differentially expressed gene lists, analyzed by DAVID. Link to high-definition figure: https://www.dropbox.com/s/l9zwx4ukw9zcs8w/Fig%201.pdf?dl=0

Next, we sought to systematically characterize gene expression profiles of NPCs in low and high CHIR conditions by mRNA-seq. Additionally, to explore the effect of low CHIR on NPCs, we generated data removing CHIR from the culture (‘No CHIR’ throughout) (Fig S1A). We also examined freshly isolated NPCs prior to culture. Low CHIR maintains expression of transcriptional regulators required for NPC specification and/or maintenance, including *Pax2*, *Wt1*, *Hoxa/d11*, *Sall1* (Fig 1C and Table S3) (McMahon 2016). In contrast, high CHIR led to a downregulation of regulators and markers of self-renewing NPCs, including *Six2* (Self et al. 2006), *Cited1* (Mugford et al. 2009), *Osr1* (Xu, Liu, et al. 2014) and *Eya1* (Xu, Wong, et al. 2014), and a concomitant increase in expression of genes associated with induction of nephrogenesis, such as *Wnt4*, *Jag1, Lhx1*, *Pax8* and *Fgf8* (Fig 1C; Park et al., 2007). Trends in gene expression from mRNA-seq were confirmed by RT-qPCR analysis (Fig S1B).

To examine biological processes at play in different NPC culture conditions, we performed gene ontology (GO) enrichment analysis of differentially expressed genes (differential expression analysis described in Methods) with DAVID (Huang da, Sherman, and Lempicki 2009). Comparing input NPCs freshly isolated from the kidney cortex with NPC-free cortical preparations (‘NFC’ throughout), the strong enrichment for NPC-relevant GO terms (Fig 1D, top panel) was consistent with a strong enrichment of Six2+ NPCs (more than 90% of isolated cells were Six2+). When NPCs cultured in high versus low CHIR conditions were compared, a strong enrichment was observed in terms associated with Wnt signaling pathway, as expected, on CHIR-mediated NPC induction (Fig 1D, middle and bottom panels). Although primary NPCs, and their counterpart in low CHIR, showed similar expression patterns for NPC self-renewal and differentiation markers, transcriptome-wide comparison of all samples clustered primary NPCs into a distinct group from NPCs cultured in either low or high CHIR (Fig S1C). This is explained by a pronounced metabolic shift in culture where there is a strong enrichment in GO analysis in metabolic processes such as sterol biosynthesis (Fig S1D). In addition, freshly isolated NPCs showed a low-level inductive signature reflecting a co-contribution of small numbers of early induced NPCs (expression of *Fgf8*, *Wnt4*, *Lhx1*, *Heyl, Bmp4*, *Mafb*, *Podxl*, Fig 1C and Table S3). Within 24 hours, low CHIR culture stabilized an undifferentiated NPCs signature with the downregulation of induction markers and upregulation of NPC-associated genes (Fig 1C). Importantly, these data show NPEM culture establishes a more rigorous model for distinguishing uninduced versus induced NPC responses than is possible with the intrinsic heterogeneity within primary isolates of NPC populations.

### CHIR-mediated induction modifies the epigenomic profile of NPCs

To investigate the chromatin landscape regulating NPCs, we integrated chromatin accessibility through ATAC-seq analysis (Buenrostro et al. 2013) with chromatin immunoprecipitation studies examining active (H3K27ac ChIP-seq) and repressive (H3K27me3 ChIP-seq) chromatin features, RNA Pol II recruitment (RNA Pol II Ser5P ChIP-Seq) and RNA-seq expression profiling (Fig S1A). Initially, we evaluated enhancers previously validated in transgenic studies (Park et al. 2012) associated with *Six2* expression in uncommitted NPCs (Six2 distal enhancer; Six2DE) and *Wnt4* activation on NPC differentiation (Wnt4 distal enhancer; Wnt4DE). In low CHIR conditions, the Six2DE shows an open and active configuration: an ATAC-seq peak, flanked by H3K27ac peaks with Pol II engagement at the enhancer and within the gene body. In high CHIR, ATAC-seq, H3K27ac and Pol II ChIP-Seq signals were reduced, correlating with the downregulation of gene expression (Fig 2A). As predicted, the Wnt4DE displayed an opposite trend in the shift from low to high CHIR: the ATAC and H3K27ac ChIP signals increased together with increased Pol II engagement in the gene body. Surprisingly, a marked enhancer specific Pol II ChIP-seq signature was visible in low CHIR NPC maintenance conditions and reduced on initiation of active transcription in high CHIR (Fig 2B).

**Figure 2.**
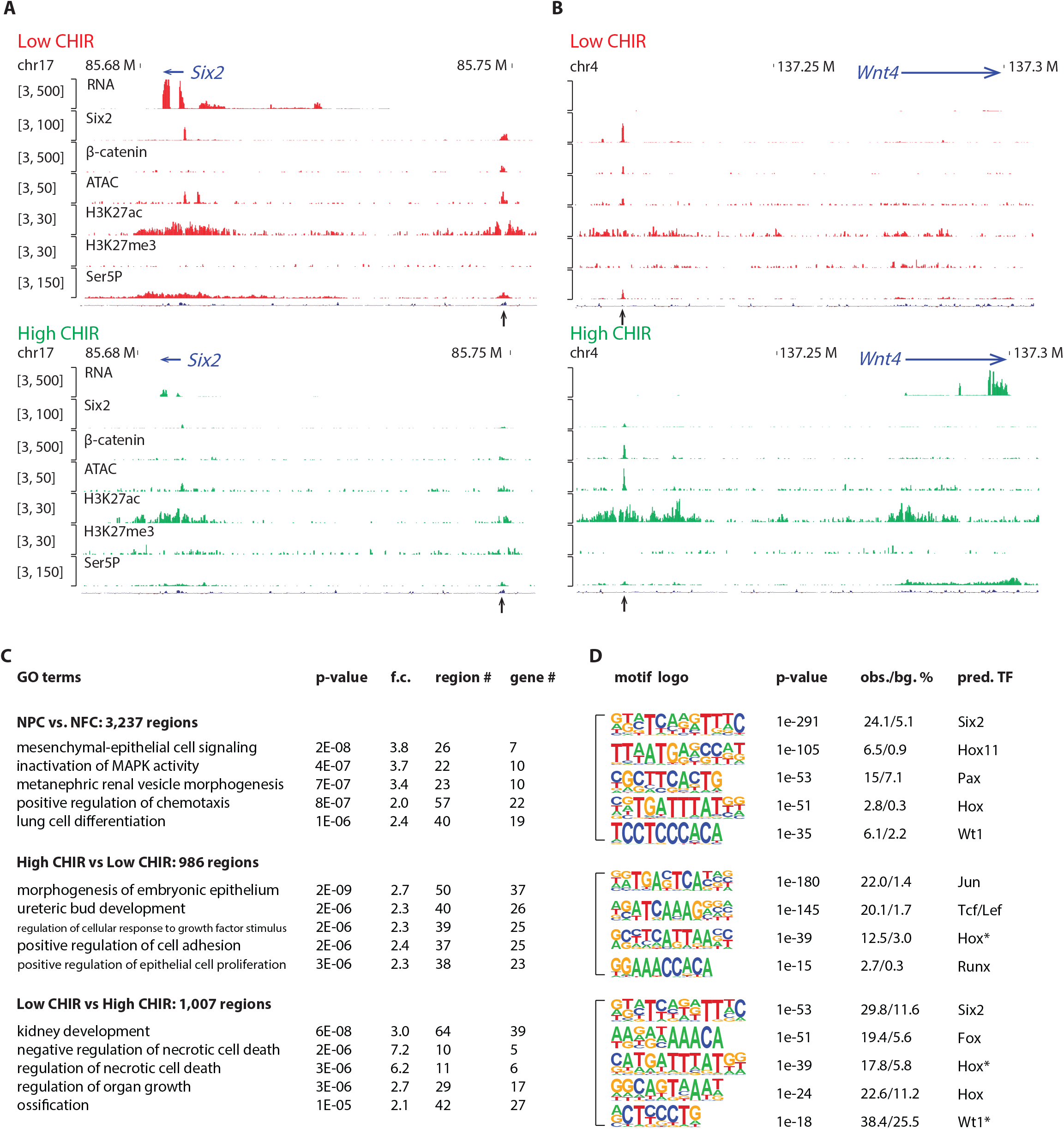
High dosage of CHIR99021 triggered change of NPC epigenome. (A and B) Genome browser view of RNA-Seq, ATAC-Seq, as well as Six2, H3K27ac, H3K27me3 and Ser5P ChIP-Seq data near *Six2* (A) and *Wnt4* (B) in low CHIR (left) and high CHIR (right) conditions. Black Arrow indicates Six2DE, Wnt4DE, respectively. (C) Top 5 most significant gene ontology terms associated with the differentially accessible (DA) regions extracted from the indicated comparisons. (D) Top 5 most enriched motifs discovered *de novo* in the indicated DA regions; * indicate less well conserved motifs for the factor.

Having validated the datasets at target loci, we examined the datasets more systematically for broad features of epigenetic regulation. We focused on the differentially accessible (DA) regions enriched in uncultured NPCs relative to NFC, as they reflect a general NPC-specific signature. The NPC-specific DA regions were significantly enriched in transcription factor binding sites for Six, Pax, Wt1 and Hox factors consistent with the critical roles of Six1/2, Pax2, Wt1 and Hox11 paralogues in NPC programs (Fig 2D) (Naiman et al. 2017, Self et al. 2006, Kreidberg et al. 1993, Wellik, Hawkes, and Capecchi 2002). Functionally, GO-GREAT analysis (McLean et al. 2010) predicted these regions were enriched near genes linked to kidney development (Fig 2C).

Hierarchical clustering of ATAC-Seq data was used to examine the relationship between CHIR dosage and the open chromatin landscape identifying differentially accessible (DA) regions in NPCs cultured in low and high CHIR conditions (Fig S2A). Most of the DA regions are distal to the transcriptional start site (TSS) of genes, indicative of enhancer elements (Fig S2B). The top two motifs identified in high CHIR-specific DA regions were the Jun (AP-1) and TCF/LEF motifs (Fig 2D), supporting a β-catenin driven increase in accessibility through engagement with TCF/LEF factors. Phosphorylated Jun has been detected in both NPCs and differentiate renal vesicles (Muthukrishnan et al. 2015). Further, Jun binds Tcf7l2 to cooperatively activate gene expression in Wnt-dependent intestinal tumors (Nateri, Spencer-Dene, and Behrens 2005). Elevated expression of *Jun*, *Junb* and *Jund* in high CHIR (Table S3) suggests a potential interplay of Jun family members with β-catenin/Tcf-driven NPC differentiation.

### Differential expression and DNA binding of TCF/LEF family members in the regulation of NPC programs

TCF/LEF factors directly bind to DNA and mediate the transcriptional response elicited by Wnt/ β-catenin. Studies in other developmental systems have generally documented repressive roles for Tcf7l1 and Tcf7l2, and activating roles for Tcf7 and Lef1 (Lien and Fuchs 2014). To examine the role of TCF/LEF factors directly in NPC maintenance and differentiation, we characterized expression of each of the four members (*Tcf7l1*, *Tcf7l2*, *Tcf7* and *Lef1*). Of the four genes, *Tcf7l1*, *Tcf7l2* and *Tcf7* transcripts were expressed at low (2-10 TPM; *Tcf7l2* and *Tcf7*) or moderate (50 TPM; *Tcf7l1*) levels in low CHIR NPC maintenance conditions. CHIR-mediated induction of NPCs resulted in a significant downregulation of *Tcf7l1* and *Tcf7l2* expression, while expression of both *Tcf7* and *Lef1* was markedly upregulated (Fig 3A and S3A-B). The same trend was observed examining the level of each protein in the nucleus of NPCs through quantitative immunofluorescence (Fig 3B-C) and Western blot (Fig S3C) analyses. To compare *in vitro* findings with NPCs *in vivo*, single cell RNA-seq transcriptional profiles were examined in cells isolated from E16.5 kidney cortex. In agreement with *in vitro* data, Tcf7l1 transcripts were enriched in self-renewing NPCs (Fig S3 F-G) while Lef1 levels were elevated in differentiated NPCs (Fig S3 F-G), though *Tcf7* and *Tcf7l2* expression levels were relatively low, with little variation between non- and early-induced NPC-states (Fig S3D-G).

**Figure 3.**
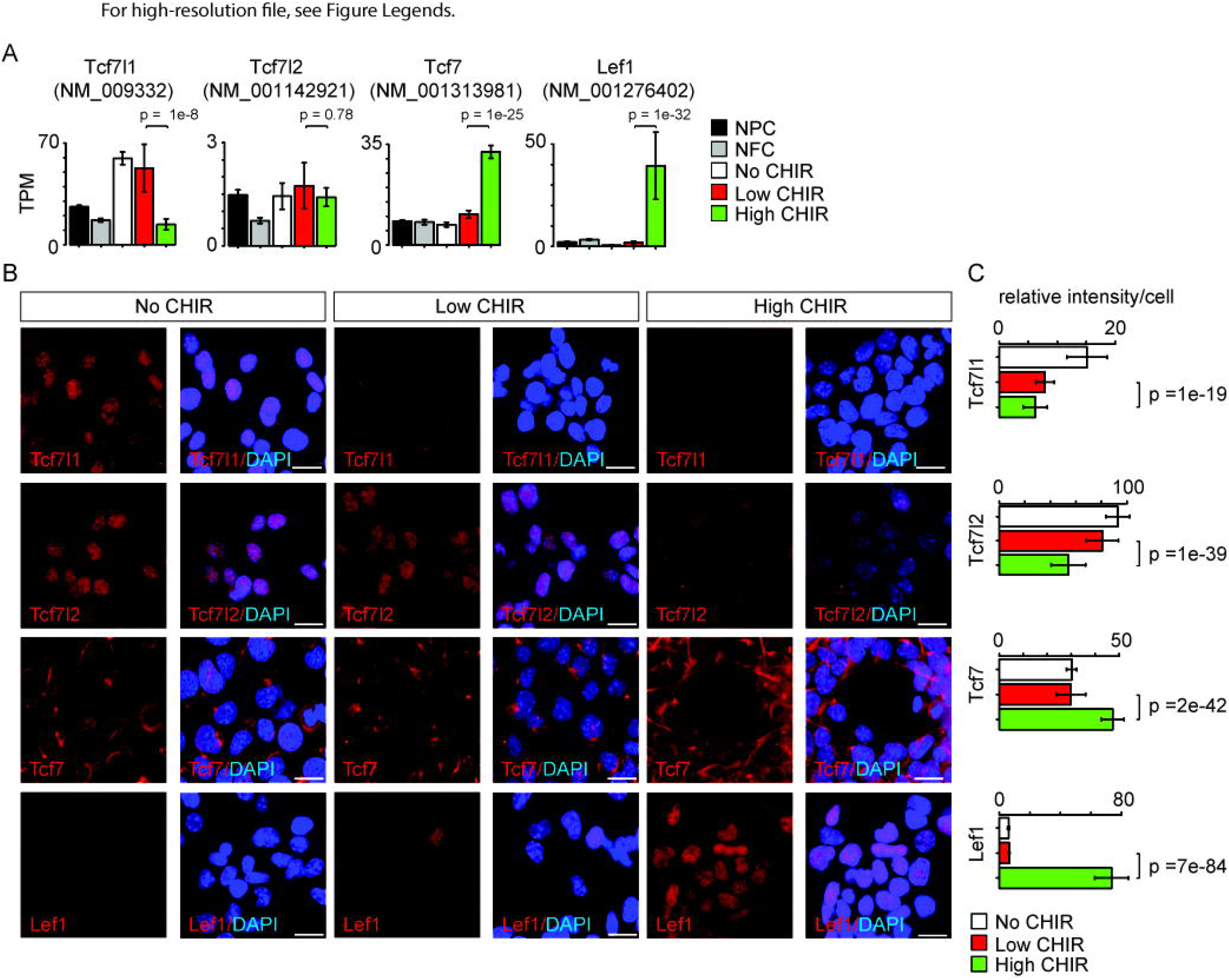
Differential expression of Tcf family transcription factors in NPC in response to distinct level of CHIR. (A) Barplots showing RNA-Seq measured expression levels of TCF/LEF family factors in NPC cultured in NPEM culture supplemented with various concentration of CHIR. (B) Immuno-fluorescence (IF) staining of TCF/LEF family factors in NPEM cultured with conditions indicated. (C) Relative intensity of IF signals from individual cells in experiment associated with B. Link to high-definition figure: https://www.dropbox.com/s/rqza0z1xpzvdrh6/Fig%203.pdf?dl=0

To directly address TCF/LEF target interactions and β-catenin-mediated regulation of NPCs, we generated Lef1, Tcf7, Tcf7l1, Tcf7l2 and β-catenin ChIP-Seq data sets from freshly isolated, uncultured NPCs, and NPCs cultured in low and high CHIR (Fig S1A). Further, given the key role for Six2 in NPC maintenance and evidence supporting Six2 engagement in Tcf7l2 and β-catenin containing complexes (Park et al. 2012), we collected Six2 ChIP-seq datasets in the same conditions. Motif discovery of TCF/LEF/β-catenin ChIP-seq binding sites shows highest DNA motif enrichment as the TCF/LEF binding element, indicating a high specificity of the data sets and supporting direct TCF/LEF/β-catenin target interactions (Fig S4C). In addition, a Hox motif is highly enriched in TCF/LEF binding sites in both low CHIR and high CHIR conditions consistent with Hox11 paralog regulation of the NPC state (Wellik, Hawkes, and Capecchi 2002, Park et al. 2012). A Runx motif is enriched in TCF/LEF binding sites in high CHIR. Runx1 expression is upregulated in the same condition (Table S3), but the significance of possible Runx regulation remains to be determined.

Consistent with the different levels of each protein in low and high CHIR conditions, Tcf7l1 engagement at DNA targets was reduced on NPC induction, while Tcf7, Lef1 and β-catenin showed a marked increase in DNA bound sites (Fig S4A). In both low and high CHIR conditions, β-catenin association overlapped extensively with the binding of cognate TCF/LEF factors specifically enriched in each condition (Fig S4B). In either CHIR condition, motif recovery suggests direct engagement through TCF/LEF binding sites (Fig S4C). However, hierarchical clustering indicates the general feature of TCF/LEF factor binding in low CHIR and high CHIR were distinct and determined by the condition as different TCF/LEF factors targeted common genomic regions in either low CHIR or high CHIR conditions (Fig S4D). This observation was most evident examining *Tcf7l2*. *Tcf7l2* mRNA and protein levels did not vary greatly between low and high CHIR conditions but Tcf7l2 DNA interactions differed significantly (Fig S4D). Our previous studies identified (Park et al. 2012) and functionally validated (O’Brien et al. 2018) a Wnt4 distal enhancer (Wnt4DE, Fig 2B) driving *Wnt4* expression in response to NPC induction. This enhancer was shown to interact with Six2, Hoxd11, Osr1 and Wt1, critical determinants of the NPC state (O’Brien et al., 2018). Our data demonstrates Tcf7l1 and Tcf7l2 were bound to the Wnt4DE in low CHIR condition, but replaced by Tcf7 and Lef1 in high CHIR conditions (Fig 4A). This switch in TCF/LEF factor binding correlated with activation of *Wnt4* expression, consistent with a potential repressive role for Tcf7l1/Tcf7l2 interactions and activating role for Tcf7/Lef1 engagement (Fig 4A). Interestingly, β-catenin was engaged at the Wnt4DE in low CHIR condition where *Wnt4* was transcriptionally silent, though β-catenin binding at this enhancer was increased on high CHIR induction of NPCs (Fig 4A). In summary, though elevating β-catenin levels through high CHIR stabilization leads to some increase in binding of β-catenin at the *Wnt4* enhancer, a marked switch in the binding signature of TCF/LEF factors is a more striking correlation with subsequent activation of *Wnt4* transcription.

**Figure 4.**
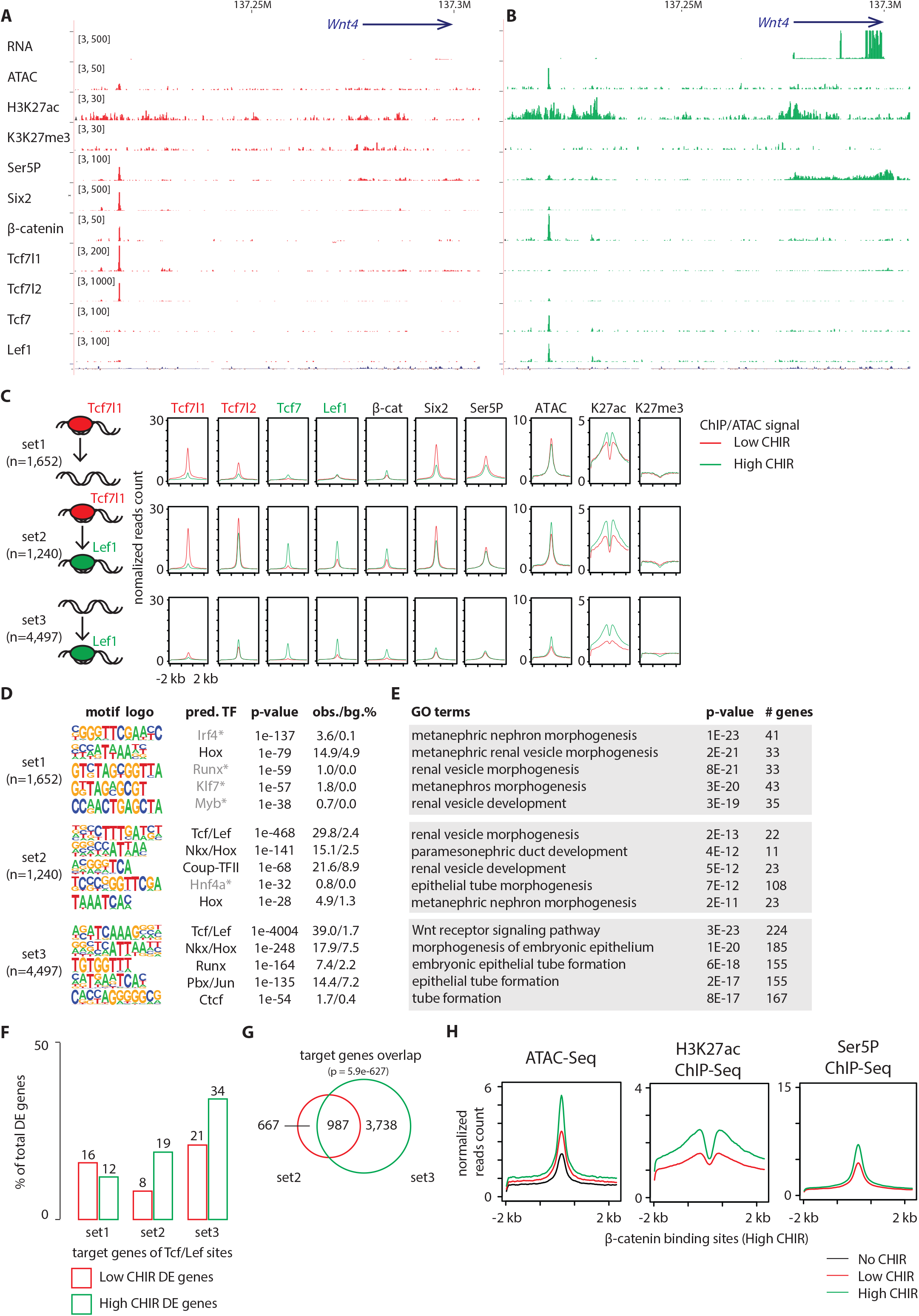
Increased CHIR dosage induces a switch of TCF/LEF factors binding to the genome. (A and B) Genome browser view of Wnt4 enhancer locus showing ChIP-Seq signal of TCF/LEF factors in low and high CHIR conditions. (C) Histograms showing binding intensity of TCF/LEF factors and chromatin markers on the 3 sets of TCF/LEF binding sites assigned by overlap of low CHIR Tcf7l1 and high CHIR Lef1 binding sites. (D) Result from de novo motif discovery of the 3 sets of TCF/LEF binding sites described in A. (E) Top Gene Ontology terms associated with the corresponding sets of TCF/LEF binding sites shown in B. (F) Percentage of TCF/LEF target genes belonging to different sets in differential expressed genes specific to low CHIR or high CHIR conditions as described in Fig. 1. (G) Venn diagram showing overlap of set 2 and 3 target genes assigned by GREAT (McLean et al. 2010). (H) Histograms showing quantification of reads from the indicated data sets in +/−2kb of β-catenin binding sites in high CHIR condition.

As Tcf7l1 and Lef1 binding best distinguished distinct NPCs’ states, we performed a more extensive analysis of these factors. Comparing DNA regions bound by Tcf7l1 in low CHIR and those bound by Lef1 in high CHIR, we observed a significant overlap (p value = 1e-569), though over half were unique to a dataset (Fig 4C). We assigned the Tcf7l1/low CHIR-specific sites as set 1 (also ‘lost,’ as no longer bound by TCF/LEF in high CHIR condition), overlapping sites as set 2 (also ‘switch,’ as TCF/LEF factors interchange at these sites between low and high CHIR conditions) and Lef1/high CHIR-specific sites as set 3 (also ‘*de novo*,’ as TCF/LEF binding sites arose on elevated CHIR levels). Interestingly, examining motif recovery across sets 1-3 indicated only sets 2 and 3 showed a strong prediction for direct TCF/LEF binding (Fig 4D). Thus, Tcf7l1 likely associates with a large number of DNA regions without direct DNA binding at TCF/LEF response elements. The majority (73%) of Tcf7l1 sites bound in low CHIR with a predicted TCF/LEF motif overlapped with those bound by Lef1 in high CHIR. In contrast, only 29% of Tcfl71 binding sites without a TCF/LEF motif prediction overlapped with Lef1 bound regions (Fig S5A). Thus, switching from a Tcf7l1 to a Lef1-centered DNA interaction at TCF/LEF response elements was a general feature of NPC induction response. Comparing with set 3 (‘*de novo*’), set 2 (‘switch’) sites displayed stronger binding of Tcf7l2, Tcf7, Lef1 and β-catenin, as well as higher level of markers of activated chromatin (ATAC-Seq and H3K27ac ChIP-Seq) in high CHIR condition (Fig 4C). Thus, the data suggests TCF/LEF sites occupied by Tcf7l1 in low CHIR are poised for stronger binding (and therefore activation) by TCF/LEF activators in high CHIR condition. Interestingly, GO GREAT analysis of all 3 sets predicted expected kidney terms such as ‘metanephric nephron morphogenesis’ or ‘renal vesicle morphogenesis’ (Fig 4E), consistent with biologically relevant interactions, arguing against artefactual responses to culture or CHIR treatment. More than half of the potential target genes of set 2 ‘switch’ sites overlaps with those of set 3 ‘*de novo*’ sites (60%; Fig 4G, p = 5.9e-627), although the latter are much broader. For this overlapping group, pre-engagement by Tcf7l1 may increase the likelihood of additional engagement in neighboring regions of DNA by Lef1, and potentially reinforce transcriptional input into a common gene target..

Examining chromatin features and RNA Pol II engagement, we observed stronger enrichment of active enhancer markers H3K27ac and RNA Pol II in low CHIR on set 1 sites predicting indirect Tcf7l1 engagement than set 2 sites where Tcf7l1 was predicted to bind DNA directly through TCF/LEF motifs (Fig 4D). Further, set 1 Tcf7l1 associated sites also showed stronger enrichment for ATAC-Seq, H3K27ac and RNA Pol II signatures. Thus, direct binding of TCF7L1/TCF7L2 on set 2 sites was associated with putative enhancers in a less active chromatin state. Consistent with this view, a direct analysis of the expression of this target gene set showed the highest expression in high CHIR conditions (Fig 4F). To examine whether elevated β-catenin correlates with chromatin activation, we examined change of active chromatin markers (ATAC-Seq and H3K27ac ChIP-Seq) and RNA Pol II loading on β-catenin bound sites. A dramatic elevation of all these signals (Fig 4H) is consistent with β-catenin engagement enhancing an open chromatin, active enhancer signature. Together, the data support the conclusion that high CHIR increased β-catenin association, correlating with switching of TCF/LEF factors at existing active chromatin, *de novo* opening of new chromatin sites, and an active transcription program initiating nephron formation.

### β-catenin uses both pre-established and *de novo* enhancer-promoter loops to drive NPC differentiation program

To examine genome organization and chromatin interactions at a higher level in self-renewing and induced NPC, Hi-C analysis was performed to characterize global chromatin-chromatin interactions (Rao et al. 2014). Further, as CTCF is known to mediate long-range chromatin interactions common across cell types, we integrated a published CTCF ChIP-seq dataset generated from NPCs isolated directly from the developing kidney (O’Brien et al. 2018).

Analysis of two Hi-C data sets, using HiCCUPS within Juicer Tools (Durand et al. 2016), replicated 19,494 low CHIR, and 20,729 high CHIR, chromatin-chromatin interaction loops (Fig. S7A). Almost half of all loops (40% low CHIR; 44%, high CHIR), including those anchored on TSSs (35% low CHIR; 49% high CHIR), were unique to each culture condition (Fig 5B). A higher number of low CHIR-specific loops associated with genes that were more highly expressed in low CHIR versus high CHIR conditions. Those loops shared between conditions and specific to high CHIR showed the reverse trend (Fig. S7B). Given a focus on chromatin loops responding to β-catenin-mediated induction of NPCs, we examined loops anchored on β-catenin peaks identified in high CHIR conditions. Of the 5,530 β-catenin associated sites in high CHIR condition, 28% (1,573) were located in loop anchors; 41% (647) of these connected to a TSS. Of the 647 peaks looped to TSSs, 57% (371) were connected by ‘conserved’ loops, shared between low CHIR and high CHIR conditions, the remainder appeared *de novo* under high CHIR conditions (Fig 5A).

**Figure 5.**
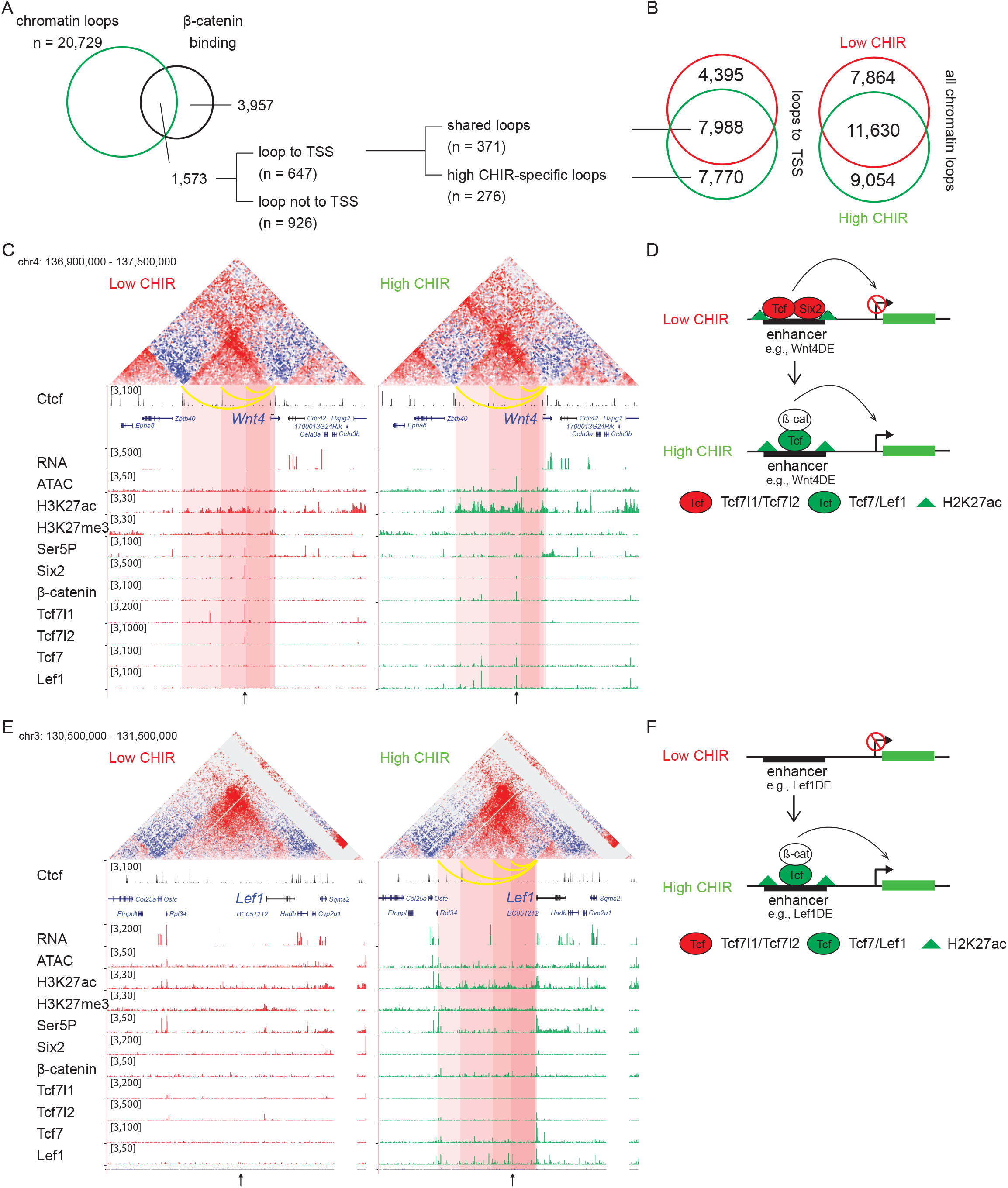
β-catenin activates gene expression through both stable and *de novo* enhancer-promoter loops. (A) β-catenin binding sites that overlap with an anchor of chromatin loop in high CHIR, the proportion that connects to a TSS (grey in the pie chart) and segregation between 2 types of loops defined in B. (B) Overlap of chromatin loops between low CHIR and high CHIR conditions. (C, D, E, F) Examples (C, E) and schematics (D, F) of β-catenin utilizing low/high CHIR-shared enhancer-promoter loops to activate Wnt4 (C, D) or high CHIR-specific loops to activate Lef1 (E, F). Black arrow at the bottom indicates the β-catenin binding sites involved in the loops.

To understand the biological consequences of these regulatory events, we identified promoters connected to β-catenin-bound enhancers. Among those connected by conserved loops present in both CHIR conditions, 56 genes were highly expressed in high CHIR conditions (Table S4), including *Wnt4*, *Lhx1*, *Emx2*, *Bmp7* and *Cxcr4* which associate with NPC differentiation *in vivo*. Of these, only *Wnt4* enhancer elements have been rigorously defined through transgenic studies (O’Brien et al. 2018). In low CHIR conditions, Six2, Tcf7l1, Tcf7l2 and β-catenin bind to the *Wnt4* distal enhancer (Wnt4DE) while in high CHIR, Tcf7 and Lef1 replace Tcf7 and Tcf7l2, with a concomitant decrease in Six2 and increase in β-catenin association (Fig 5C). Two additional loops further upstream anchor to Ctcf binding sites and loop to the *Wnt4* TSS (Fig 5C). Interestingly, all three loops are stable between low and high CHIR conditions consistent with a pre-determined chromatin organization facilitating rapid activation of a nephron forming inductive program, on switching the TCF/LEF transcriptional input (Fig 5D).

*Lef1* was amongst the gene sets defined by highly enriched expression in high CHIR conditions with *de novo* loop formation in high CHIR (Fig 5F and Table S4). The *Lef1* locus showed no loop connections to the *Lef1* TSS in low CHIR. Further, no TCF/LEF/β-catenin binding was detected, and ATAC-seq and H3K27 acetylation showed only background levels around the *Lef1* locus. In contrast, four interaction loops appeared in high CHIR conditions, one of which associated with a strong Lef1/β-catenin interaction site, two others mapped to CTCF bound regions (Fig 5F). In high CHIR, a string of LEF1/TCF7/β-catenin binding events accompanied enhanced accessibility (ATAC-seq) and the appearance of an active chromatin signature (H3K27ac) (Fig 5F). These data suggest *de novo* loop formation may follow from *de novo* interaction of LEF1/TCF7/β-catenin binding complexes at the *Lef1* enhancer, in a feedback mechanism driving NPC commitment (Fig 5F).

## Discussion

Genetic analysis of mouse models point to a requirement for β-catenin in both the maintenance (Karner et al. 2011) and differentiation (Park, Valerius, and McMahon 2007) of NPCs. Previous studies directly examining β-catenin association in differentiating NPCs showed direct engagement of β-catenin at enhancers regulating expression of differentiation promoting genes such as *Wnt4* (Park et al. 2012). Data here confirmed these earlier findings and extended our understanding through a comprehensive analysis of Wnt-directed transcriptional engagement and epigenetic organization, in a simple *in vitro* model of mammalian NPC programs. TCF/LEF factors are the transcription factors that ultimately mediate the transcriptional response to Wnt signaling. The direct analysis of all four TCF/LEF factors enables several key observations to be made, and conclusions drawn, from the analysis of NPC responses to CHIR modulation of β-catenin levels.

1) In low CHIR, NPC maintenance and expansion conditions, the mitogenic activity mediated through CHIR stimulation of Wnt-signaling appears to be independent of TCF/LEF binding to TCF/LEF motifs in target genes; 2) CHIR-mediated elevation of β-catenin is accompanied by reduced expression of mRNAs for transcriptional inhibitory TCF factors (Tcf7l1 and Tcf7l2), and dramatic increase in expression of mRNAs for activating forms (Tcf7 and Lef1), promoting an inductive program; 3) Direct binding of inhibitory Tcf7l1 and Tcf7l2 engagement at target motifs within putative cis-regulatory elements prefigures engagement of Tcf7 and Lef1 in a chromatin landscape primed for the transcriptional activation of the nephron forming program; 4) High CHIR invokes a switch from inhibitory to activating TCF/LEF engagement at enhancers promoting nephron differentiation consistent with β-catenin controlling TCF/LEF target engagement. 5) In addition to enhancer-promoter loops pre-established in uncommitted NPCs associated with switching of inhibitory to activating TCF/LEF binding signatures on high CHIR induction, TCF/LEF/ β-catenin interactions at *de novo* sites may play additional roles in the inductive process, including positive feedback in the Lef1 program promoting nephrogenesis.

### Wnt signaling, β-catenin and TCF/LEF factors in NPC maintenance and expansion

Though genetic evidence supports a direct role for β-catenin in regulating the maintenance and expansion of NPCs, and low CHIR activity is essential for normal NPC expansion *in vitro*, direct analysis of Tcf, Lef and β-catenin engagement does not provide support for a direct transcriptional mechanism mediated though direct TCF/LEF DNA interactions with low CHIR-dependent target genes. We directly examined 16 genes identified in genetic screens *in vivo* to display *Wnt9b-* dependent expression (Karner et al. 2011). Seven of the sixteen showed elevated expression of at least one isoform in low CHIR versus no CHIR (Fig. S1G and Table S5) consistent with a Wnt-signaling input. Two of this group, *Pla2g7* and *Tafa5/Fam19a5,* wee also reported to be ectopically activated by LiCl stimulation of the Wnt pathway and ectopic activation of β-catenin (Karner et al., 2011). Our data corroborated the up-regulation of Tafa5/*Fam19a5* but not *Pla2g7* in high CHIR (; Fig. S1G and Table S5). However, ChIP-qPCR data arguing for β-catenin engagement around the TSS bindings sites of *Pla2g7* and *Fam19a5/Tafa5* could only be corroborated near one site in our datasets (Fig. S6). Comparison of low CHIR and no CHIR GO enrichment analysis revealed highly significant enrichment in cell cycle-related terms (Fig. S1E), consistent with pro-proliferation roles of β-catenin in self-renewing NPCs. However, among the 85 genes associated with the term ‘cell cycle’, only one gene shown significant β-catenin association within 500 kb of the TSS in low CHIR (Fig. S1F), arguing against a scenario that direct transcriptional regulation of these genes through β-catenin engagement.

These findings raise the possibility that β-catenin acts through an alternative transcriptional mechanism. A prominent association of TCF7L1 is observed in low CHIR to DNA regions where the absence of a TCF/LEF motif may suggest indirect means of association, through protein-protein interactions. However, no strong consensus target emerges from examining motif enrichment in this subset of the TCF7L1 binding data (Fig. 4D). Alternatively, β-catenin may play an essential, non-transcriptional role that links to control of cell proliferation. TCF-β-catenin nuclear complexes have been reported to oscillate with the cell cycle suggesting potential nuclear roles independent of DNA association (Ding et al., 2014). Further, β-catenin is reported to play a essential, non-transcriptional role in self-renewal of mouse epiblast stem cells (Kim et al. 2013). Importantly, our studies cannot exclude unknown complicating actions of CHIR-mediated inhibition of GSK3 outside of β-catenin regulation or GSK3-independent CHIR-responses. Wnt-ligand mimetics offer a promising future approach to confirm CHIR-β-catenin-centered findings in the current study.

In mouse embryonic stem cell (mESC) culture, canonical Wnt signaling, induced by CHIR99021, supports long-term self-renewal of ESCs (Ying et al. 2008). Tcf7l1 has been shown to repress expression of genes involved in stem cell maintenance while Tcf7 and Lef1 activate targets (Yi et al. 2011, Wray et al. 2011) consistent with a classic canonical Wnt transcriptional activation program of stem cell renewal. However, other studies directly analyzing β-catenin interactions at the chromatin level suggest an indirect process where β-catenin may block the negative interplay of Tcf7l1 binding at the Sox motif of Sox-Oct bound stem cell promoting enhancers (Zhang et al., 2013). In this ESC system, β-catenin target sites containing TCF/LEF motif correlate strongly with differentiation promoting targets, consistent with the normal role of β-catenin *in vivo* in regulating gastrulation (Haegel et al. 1995). Addition of a second small molecule (PD03) inhibiting MEK/ERK signaling is essential to block this differentiation promoting activity (Zhang et al. 2013). Further, depletion of both Tcf7l1 and Tcf7 are sufficient to maintain Wnt ligand-independent expansion of ESCs, consistent with the implication that β-catenin activation can abrogate the repressive effect of TCF7L1 independent of an activator TCF7 (Yi et al. 2011).

In hair follicle stem cells (HFSC), Wnt induces a transition of the stem cells from the quiescent to the proliferative state. Tcf7l1 and Tcf7l2 are preferentially expressed in HFSC, while Lef1 and Tcf7 are preferentially expressed in the differentiated HFSC, i.e. hair germ (HG) cells (Merrill et al. 2001, Lien et al. 2014). TCF7L1/TCF7L2 repress genes involved in HFSC differentiation, which are activated by Tcf7/Lef1, correlating with replacement of TCF7L1/TCF7L2 by Tcf7/Lef1 on relevant enhancers, a close parallel to activity in NPC programs described here (Adam et al. 2018).

### Elevating β-catenin leads to activation of Tcf/Lef-bound enhancers

In the Wnt-off state, Tcf factors are known to recruit Groucho family co-repressors (Cavallo et al. 1998) and histone deacetylase (Billin, Thirlwell, and Ayer 2000) to repress Wnt target gene expression.. Upon Wnt ligand stimulation, Groucho is replaced by β-catenin for activation (Brantjes et al. 2001, Daniels and Weis 2005). From evidence *in vitro*, β-catenin has been shown to be able to recruit various chromatin modulators, including histone acetyl transferase (HAT) (Hecht et al. 2000), histone methyl transferase (Sierra et al. 2006), and chromatin remodeler (Barker et al. 2001). Furthermore, through interaction with Pygo and Bcl9 (Kramps et al. 2002, Schwab et al. 2007), as well as direct interaction (Kim et al. 2006), β-catenin can form a complex with the Mediator complex, which bridges the Tcf-bound enhancer to RNA Pol II complex at the target gene promoter (Jeronimo and Robert 2017). In addition, Six2 binding at certain TCF/LEF targets in NPCs might also confer a repressive effect; Six2 interacts with histone deacetylation complexes (HDACs), and depletion of the HDACs in NPC elevates *Wnt4* and *Lef1* expression (Liu et al. 2018). Similarly, the repressive chromatin modifier *Ezh1* and *Ezh2* also repress Wnt4 and Lef1 expression in NPC through maintaining H3K27me3 mark (Liu et al. 2020), although the repressive functions of such chromatin modifiers tend to have broader effect

The potential for an extended interaction of Tcf7l1 and Tcf7l2 with co-repressors in suppressing the NPC commitment program has not been addressed in this study.. However, given the observation that activation of target genes committing NPCs to a nephrogenic program correlates with a switch to Tcf7 and Lef1 engagement, it seems unlikely removal of a co-repressor input would be sufficient for full activation in the absence of these strong activators. The observed shift in TCF/LEF factor engagement at DNA targets through elevating CHIR levels raises the question of how rising levels of β-catenin might regulate this transcriptional switch. Given a dual role for Tcf7l2 as both a transcriptional repressor and activator in canonical Wnt transcription (Korinek et al. 1997, Chodaparambil et al. 2014, Lien and Fuchs 2014), β-catenin may switch Tcf7l2 to an activator state. Alternatively, low levels of Tcf7 present in low CHIR may be sufficient for β-catenin engagement and transcriptional activation. In this model, binding of a Tcf7-β-catenin complex would be favored over inhibitory TCF complexes. In either scenario, it is likely that the transcriptional up-regulation of Tcf7 and Lef1 creates a feed-forward loop to amplify the transcriptional activation response. Distinguishing amongst these possibilities will require effective and sensitive strategies to specifically modify regulatory components in the experimental model system.

### Pre-establishment of enhancer-promoter loops prefigures a nephrogenic program

Classical embryological studies have identified two broad categories of inductive processes by which uncommitted stem or progenitor cells make subsequent cell fate choices (Saxén and Sariola 1987). Instructive signaling leads to cells adopting distinct cell fates each determined by the signaling input. In this scenario, cells have multiple fate choices open to that cell. In contrast, permissive signaling only leads to a single outcome. The observation that nephron anlagen only undergo a restricted nephrogenic response, and no other, to inductive signals, was taken as evidence that nephron progenitor cells are in an inflexible regulatory state, predetermined for kidney formation. Studies in *Drosophila* have indicated that during development, certain enhancer-promoter loops are stable (Ghavi-Helm et al. 2014), i.e. the enhancer-promoter loop is established before the target gene is activated as a mechanism to prime developmentally potent cells to differentiate into pre-destined cell fates.

Analysis of Hi-C data shows enhancer-promoter loops present in low CHIR condition are consistent with NPCs exhibiting a primed genomic state promoting nephron forming programs. Approximately 56% of loops observed in high CHIR were observed in low CHIR. Of these, 70% connected to a TSS (Fig 5B). Example of genes where such ‘conserved’ loops connect TCF/LEF/β-catenin binding events to TSS include a number of genes within the nephrogenic program including Wnt4, Lhx1, Emx2, Bmp7 and Cxcr4. These data are consistent with the concept that at least part of Wnt/β-catenin activated differentiated program in NPC is primed through enhancer-TSS loop establishment, most likely at the time of specification of the NPC lineage though these remain to be determined. These findings also raise an interesting possibility that Tcf7l1 and Tcf7l2 engagement at such enhancer-promoter regions may maintain the primed state during an extensive period of progenitor expansion, in the course of kidney development. Interestingly, in low CHIR maintenance conditions, the Wnt4DE showed higher levels of PolII association than in high CHIR conditions where *Wnt4* is transcribed, consistent with a stable enhancer/promoter/PolII association in the primed state. Indeed, Tcf7l1and Tcf7l2, might serve as a platform for assembly of the transcriptional machinery to facilitate target gene activation with a sufficient level of β-catenin. Hi-C studies also identify new loop interactions consistent with *de novo* gene activation, notably in a predicted Lef1 feed-forward loop..

In summary, NPC culture provides a powerful model for deepening a mechanistic understanding of the regulatory processes balancing maintenance, expansion, and commitment of NPCs. Bulk isolation of stem progenitor cells is problematic for many stem/progenitor systems. Further, *in vitro* conditions enabling the controlled switching been stem/progenitor and differentiation programs have only been described for a few of these systems. Given a broad role for Wnt/β-catenin signaling in regulating stem and progenitor cell programs in metazoans, the findings here may have broader significance for Wnt-directed control of organogenesis. Further, Notch and PI3K activity can also drive early nephrogenic responses *in vivo* or *in vitro* (Lindstrom et al. 2015, Boyle et al. 2011). The NPC culture model will provide a rigorous analytical platform for future exploration of how distinct pathway activities are integrated in the induction of mammalian nephrons

## Materials and Methods

### mRNA-Seq and data analysis

50,000 – 100,000 cells were collected for each RNA experiment. RNA was isolated with RNeasy micro kit (Qiagen, #74004). mRNA-Seq libraries were prepared with KAPA Stranded mRNA-Seq Kit (Kapa Biosystems, #KK8420). The libraries were subsequently sequenced with Illumina NextSeq500 model with pair-end 75 bp setting.

mRNA-Seq reads were aligned with STAR (Dobin et al. 2013) to mm10 assembly and quantified with Partek E/M to generate a count table, and finally converted to TPM for representation. All the steps above were implemented in the Partekflow web platform (St. Louis, MO, USA) sponsored by USC Norris Medical Library.

To identify differentially expressed genes, count tables of the two groups of data being compared were processed through DESeq2 (Love, Huber, and Anders 2014) to obtain the negative binomial p values which evaluates the significance of difference by read counts. The differentially expressed genes were defined with the following threshold: TPM > 5, fold change >3, negative binomial p value < 0.05, unless otherwise specified.

Gene Ontology enrichment analysis was performed with DAVID (Huang da, Sherman, and Lempicki 2009).

### ChIP-Seq

1. Fixation. Freshly isolated nephron progenitor cells were fixed in 1 mL AutoMACS running buffers (for each 3-5 million cells). Cultured nephron progenitor cells were fixed in NPEM medium before scraping. In both cases, cells were fixed with final 1% formaldehyde (Thermo Fisher Scientific, #28908) for 20 min at room temperature.
2. Chromatin preparation. 3-5 million cells were processed for each chromatin preparation. Chromatin preparation includes cell lysis and nuclei lysis, which were done with SimpleChIP® Sonication Cell and Nuclear Lysis Buffers (Cell Signaling Technology #81804) following manufacturer’s instruction.
3. Chromatin fragmentation. For chromatin fragmentation, lysed nuclei were sonicated with Branson Ultrasonics Sonifier S-450, using a double-step microtip. Each sample was re-suspended in 1 mL nuclear lysis buffer in a 15 mL conical tube, embedded in water-filled ice. Sonication was performed at 20% amplitude for 4 min, with 3 secs of interval after each 1 sec of duty time.
4. Immuno-precipitation. 1 million-equivalent fragmented chromatin was used for each immunoprecipitation experiment. Immuno-precipitation was done with SimpleChIP® Chromatin IP Buffers (Cell Signaling Technology #14231) following the manufacturer’s instruction, with the following details. The amounts of antibody used were case-dependent. In general, 2 μg or 1:50 to 1:100 antibody was used for each precipitation. Chromatin with antibody were rotated overnight at 4 °C before 1:40 protein A/G agarose beads (Thermo Fisher Scientific, #20423) were added and incubated for another 6 hrs. to overnight. After washing and elution, antibody-precipitated input DNA were purified with minElute reaction cleanup kit (Qiagen, #28204), reconstituting to 35 uL EB buffer.
5. ChIP-qPCR. qPCR was performed with Luna® Universal qPCR Master Mix Protocol (New England Biolab #M3003) on a Roche LightCycler 96 System. For each reaction, 0.5 out of 35 uL ChIP or input DNA was used. The qPCR primers used are listed below:

Six2-DE:
F: ggcccgggatgatacatta
R: cgggtttccaatcaccatag
Wnt4-DE:
F: GACCCATAAGGCAGCATCCA
R: CTTGCTGGGCAGAGATGAA
Non-ChIP:
F: tctgtgtcccatgacgaaaa
R: ggaagtcatgtttggctggt
6. Sequencing. ChIP-Seq libraries were prepared with Thruplex DNA library prep kit (Clontech, # R400523). The libraries were sequenced with Illumina NextSeq500 model using single-end 75 bp setting.

### ChIP-Seq data analysis

ChIP-Seq reads were aligned with bowtie2. The alignment files are filtered to remove duplicate reads with Picard (http://broadinstitute.github.io/picard/index.html). Peak calling was performed with MACS2 (Feng et al. 2012) with combined replicate data sets of the ChIP/condition being considered, and using combined replicate input from the same condition as control. To obtain relatively strong peaks, the peaks were first filtered for q-value < 1e-4. Afterwards, the counts of normalized reads were generated within +/− 250 bp windows of the filtered peaks. To obtain consistent peaks in both replicates, we filtered the ones with > 10-fold enrichment in the +/− 250 bp window in both replicates for downstream analysis.

For data shown in Figure S4, the peaks were further filtered for those with fold enrichment > 20 in order to focus on strong peaks. Overlapping peaks were defined as those within 150 bp from each other’s center.

For visualization, wiggle tracks were generated with QuEST (Valouev et al. 2008). The intensity of peaks is measured as fold enrichment, which is calculated by the number of reads within the +/− 250 bp window divided by the total mapped reads in the library, normalized to the size of genome. *De novo* Motif discovery and motif scan was performed with Homer (Heinz et al. 2010). Gene Ontology analysis was performed with GREAT (McLean et al. 2010). To determine the overlap of ChIP-Seq peaks, peak centers from the two compared data sets were overlapped, and centers beyond 150 bp from each other were considered as unique sites (Fig S4A-B).

### ATAC-Seq and data analysis

Each ATAC-Seq experiment was performed with 50,000 cells, following the published protocol (Buenrostro et al. 2013). ATAC-Seq libraries were sequenced with Illumina NextSeq500 model using single-end 75 bp setting.

ATAC-Seq reads were aligned with bowtie2. Peak calling was performed with MACS2 (Feng et al. 2012) without control data. Subsequently, reads from each replicate were counted with Homer within +/− 250 bp of peak center. Peak intensity was represented as fold enrichment as described in ChIP-Seq data analysis. To obtain consistent peaks, only the ones passing threshold (fold enrichment > 3) in all 3 replicates were retained for downstream analysis.

To identify enriched regions in condition A over condition B, ATAC-Seq peak coordinates from condition A were used to count reads within a +/− 250 bp windows, from both condition A and condition B. Fold enrichment in each replicate was calculated and went through DESeq2 procedure in order to identify statistically significantly differentiated accessible (DA) regions. The threshold for identifying DA regions is fold enrichment > 5, fold change > 2 and negative binomial p-value < 0.05.

To perform genome-wide hierarchical clustering, peaks from all replicate data sets in comparison were merged (peaks < 150 bp from each other are combined into one peak taking the midpoint as the new coordinate). Subsequently, ATAC-Seq reads from all samples concerned were counted in +/− 250 bp bins centering on the merged peaks, generating a count table. Hierarchical clustering was generated based on fold enrichment calculated from the count table.

*De novo* Motif discovery was performed with Homer (Heinz et al., 2010). Gene Ontology analysis was performed with GREAT (McLean et al., 2010).

### Hi-C data generation and analysis

We generated about 700 million raw reads for each sample. The reads were aligned by bwa (Li and Durbin 2009), then duplicates were removed with Picard. The Hi-C files were created and loop calling was done with the Juicer Tools (Durand et al. 2016).

To identify loops that are consistently present in both replicates, we extracted loops whose coordinates of anchors are within 10 kb between replicates.

| Condition | replicate1 | replicate2 | overlap |
| --- | --- | --- | --- |
| NPC-1.25 | 35854 | 39693 | 12181 |
| NPC-5 | 36731 | 40595 | 12231 |

To find TSS of genes and peaks connected by loops, we look for peaks that are within 5 kb from center of one of the loop anchors and TSS that are within 15 kb from center of the other loop anchor.

### Single-cell RNA-Seq and data analysis

Cortical cells were dissociated from E16.5 kidneys as described in Brown et al., 2015. Single-cell RNA-Seq library was synthesized using the 10X Genomics Chromium platform, with v2 chemistry and reads were mapped with Cell Ranger, as described in Lindstrom et al (Lindstrom, Guo, et al. 2018). Unsupervised clustering of transcriptional profiles, feature plots and dot plots were generated with Seurat v2.3 (Satija et al. 2015). To computationally isolate the nephron lineage, we subset clusters of cells for those highly expressing nephron lineage markers (Six2, Wnt4, Wt1) but not interstitial progenitor cells markers (Foxd1, Meis1, Fat4).

### Reverse transcription followed by qPCR (RT-qPCR)

Total RNA was reverse-transcribed with SuperScript™ IV VILO™ Master Mix with ezDNase™ Enzyme

(cat #: 11766050). qPCR was performed with Luna® Universal qPCR Master Mix Protocol (New England Biolab #M3003) on a Roche LightCycler 96 System. p-values were obtained by performing t-test between replicates of samples indicated. Primers used in RT-qPCR are listed as follows:

Six2:
F: CACCTCCACAAGAATGAAAGCG
R: CTCCGCCTCGATGTAGTGC
Cited1:
F: AACCTTGGAGTGAAGGATCGC
R: GTAGGAGAGCCTATTGGAGATGT
Wnt4:
F: AGACGTGCGAGAAACTCAAAG
R: GGAACTGGTATTGGCACTCCT
Jag1:
F: CCTCGGGTCAGTTTGAGCTG
R: CCTTGAGGCACACTTTGAAGTA
Fgf8:
F: CCGAGGAGGGATCTAAGGAAC
R: CTTCCAAAAGTATCGGTCTCCAC
Lhx1:
F: CCCATCCTGGACCGTTTCC
R: CGCTTGGAGAGATGCCCTG
Pax8:
F: ATGCCTCACAACTCGATCAGA
R: ATGCGTTGACGTACAACTTCT
Tcf7l1:
F: CCCGCTGACACCTCTCATC
R: ACAGTGGGTAATACGGTGACAG
Tcf7l2:
F: AACGAACACAGCGAATGTTTCC
R: CACCTTGTATGTAGCGAACGC
Tcf7:
F: AACTGGCCCGCAAGGAAAG
R: CTCCGGGTAAGTACCGAATGC
Lef1:
F: TGTTTATCCCATCACGGGTGG
R: CATGGAAGTGTCGCCTGACAG

### Immunofluorescence staining

To perform immunofluorescence staining, cell cultures were fixed with 4% PFA in PBS for 10 min, then washed with PBS twice before blocking in 1.5% SEA block (ThermoFisher, 107452659) in TBST (0.1% Tween-20 in TBS). After minimally 30 min at room temperature, switched to primary antibody (diluted in blocking reagent) incubation in 4 degree overnight. After washing 3 times with TBST, switched to secondary antibody (diluted in blocking reagent) incubation for minimally 45 min in room temperature, blocking light. This was followed by 3 washes with TBST, then the cells were kept in PBS for confocal imaging.

### Immunoblots

To separate, protein samples were boiled with β-mercaptol and ran in SDS-PAGE gels casted from 30% Acrylamide/Bis solution 29:1 (Bio-rad, 1610156) using the Mini-PROTEAN® system (Bio-rad). Afterwards, the gel was transferred in Mini Trans-Blot® Cell (Bio-rad) system to PVDF membranes (Immobilon-P, EMD Millipore, IPVH08100). The membrane with protein was blocked with I-block (Applied Biosystem, T2015) in TBST (0.1% TritonX-100 in TBS) at room temperature for 45 min before switching to primary antibody (diluted in blocking reagent) incubation in 4 degree overnight. Subsequently, the membrane was washed 3 times and switched to secondary antibody incubation (diluted in blocking reagent) for 45 min at room temperature. This was followed by 3 washes with TBST before drying the membrane and adding HRP substrate (Pierce™ ECL Plus Western Blotting Substrate, Thermo, 32132). Finally, the membrane was used on Autoradiography Film (5×7, Blue Devil, Premium, 100 Sheets/Unit, Genesee Scientific/Amazon) to visualize location of protein.

## Figure Legends

**Figure S1. Supplementary RNA-Seq data analysis.** (A) Overview of experiment design and data available (grey). (B) Barplots show RT-qPCR measurement of relative expression of the indicated genes, as verification of results in Fig. 1C. (C) Hierachical cluster of R-square values between transcriptome-wide TPM of the indicated pair of replicate RNA-Seq data sets. (D) Top 5 enriched GO terms of genes differentially expressed between low CHIR condition and uncultured NPC. (E) Top 5 enriched GO terms of genes differentially expressed between low CHIR and no CHIR conditions. (F) Venn diagram shows overlap of β-catenin target genes with all genes associated the gene ontology term ‘cell cycle’ that are highly expressed in low CHIR vs. no CHIR condition. (G) Heatmap and hierarchical cluster showing log2 relative TPM (TPM divided by mean across samples) to reflect change of gene expression (at isoform level) of Karner et al., 2011 class II genes (differentially expressed low CHIR > no CHIR or high CHIR > low CHIR) in our data set. The highlighted gene is supported by genetic evidence of regulation by β-catenin in the current data.

**Figure S2. Supplementary ATAC-Seq data analysis.** (A) Hierarchical cluster of R-square values between normalized ATAC-Seq reads within merged peaks from all samples. (B) Histograms of distances from differentially accessible (DA) chromatin regions to TSS implicate a predominant enhancer feature. (C) Top 5 most significant GO terms associated with the NFC-specific DA regions. (D) Top 5 most enriched motifs discovered *de novo* in the NFC-specific DA regions.

**Figure S3. Supplementary evidence for differential expression of TCF/LEF factors.** (A) Barplots show RT-qPCR measurement of relative expression of TCF/LEF family factors, as verification for results in Fig. 3A. (B) Barplots show expression of individual transcripts of Rcf/Lef factors in our RNA-Seq data. (C) Immunoblots of TCF/LEF family factors in NPC cultured in NPEM with the indicated treatment. (D) tSNE plot displaying unbiased cluster of nephron lineage cells profiled by single-cell RNA-Seq. (E) Feature plots displaying distribution of self-renewal (red) and differentiation (green) marker genes transcripts on the tSNE plot. (F) Feature plots showing distribution of TCF/LEF factors transcripts on the tSNE plot. (G) Dotplots showing accumulated expression level of marker genes as well as TCF/LEF factor in selected clusters of cells.

**Figure S4. Supplementary ChIP-Seq data analysis.** (A) Numbers of and overlap between binding sites of the same factors between different conditions. (B) Numbers of and overlap between binding sites of different factors in the same conditions. (C) Most enriched motifs by *de novo* discovery (Homer) from binding sites of the data sets indicated. (D) Hierachical clustering of normalized read counts of ChIP-Seq data sets on merged binding sites.

**Figure S5. Analysis of Tcf7l1 binding in low CHIR.** (A) Overlap of direct and indirect Tcf7l1 binding in low CHIR with Lef1 binding in high CHIR (left) and enrichment of ChIP-Seq reads on the two types of binding events. (B) Distribution of ATAC-Seq, H3K27ac ChIP-Seq and Ser5P ChIP-Seq signals on Tcf7l1 binding sites in low CHIR with or without TCF/LEF motifs. (C) Gene Ontology analysis by GREAT on Tcf7l1 binding sites in low CHIR with or without TCF/LEF motifs. (D) Overlap of predicted target genes between Tcf7l1 binding sites in low CHIR with or without TCF/LEF motifs. (E) Histograms showing intensities of ATAC-Seq or ChIP-Seq signals around set1, 2 and 3 TCF/LEF binding sites in low CHIR and high CHIR conditions.

**Figure S6. β-catenin binding sites near (A) Pla2g7 and (B) Tafa5, two β-catenin target genes reported in Karner et al., 2011.** The ChIP-qPCR target sites were marked as red bins on the top track. The highlighted region is where our data is consistent with Karner’s.

**Figure S7. Supplementary HiC data analysis.** (A) Reproducible loops between replicates. (B) Barplots showing the percentage of loops in the corresponding category (high CHIR only, low CHIR only or share) that are linked to differentially expressed genes (DEG) that are high in low CHIR or high CHIR.

