## Supplemental Figure 1 for "A β-catenin-driven switch in TCF/LEF transcription factor binding to DNA target sites promotes commitment of mammalian nephron progenitor cells"

Fig. S1

A

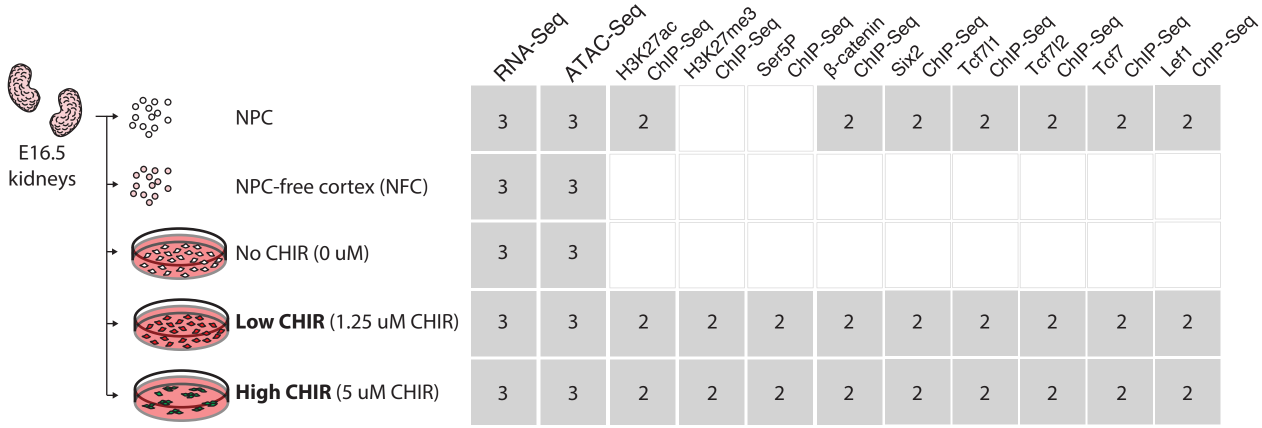


B

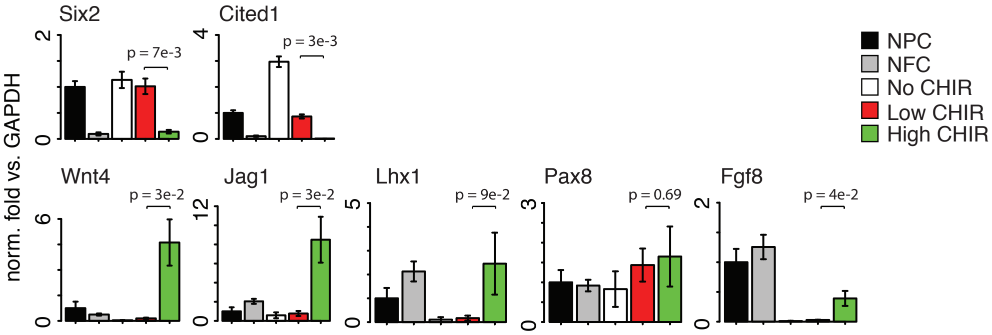


F

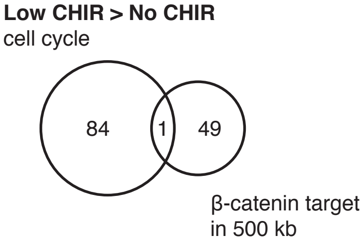


C

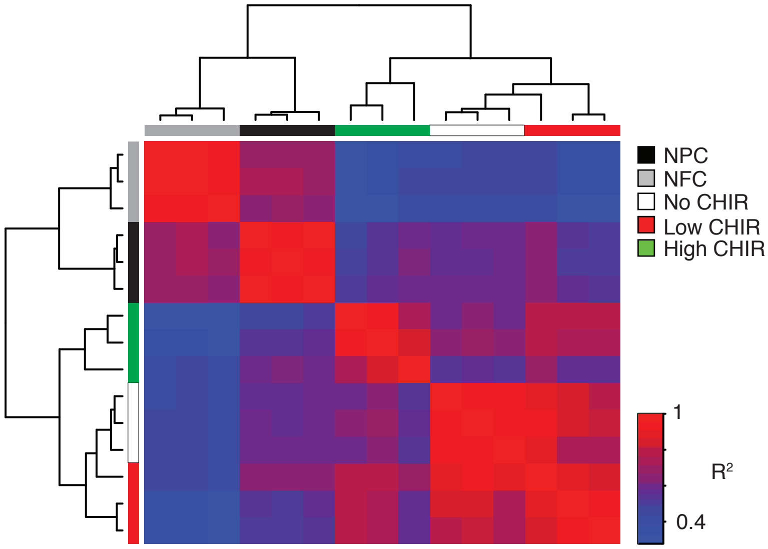


D

| GO terms | # | p-value | F.E. |
|---|---|---|---|
| **Low CHIR > NPC (494 transcripts)** | | | |
| sterol biosynthetic process | 17 | 3.7E-20 | 27.7 |
| cholesterol biosynthetic process | 17 | 1.5E-18 | 23.4 |
| steroid metabolic process | 19 | 4.2E-13 | 10.0 |
| cholesterol metabolic process | 18 | 1.4E-11 | 8.9 |
| steroid biosynthetic process | 15 | 8.2E-11 | 10.6 |
| **NPC > Low CHIR (284 transcripts)** | | | |
| response to mechanical stimulus | 11 | 7.3E-09 | 13.5 |
| multicellular organism development | 36 | 1.4E-07 | 2.7 |
| skeletal muscle cell differentiation | 9 | 4.9E-07 | 12.6 |
| positive regulation of transcription from RNA polymerase II promoter | 33 | 1.7E-06 | 2.6 |
| positive regulation of transcription, DNA-templated | 23 | 5.8E-06 | 3.1 |

G

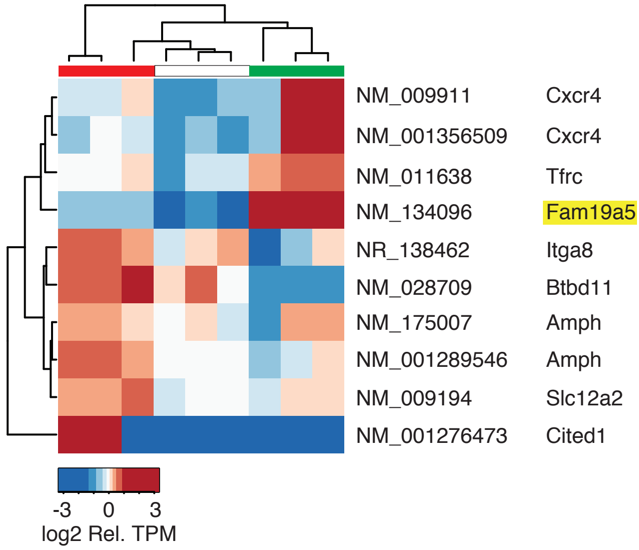


E

| GO terms | # | p-value | F.E. |
|---|---|---|---|
| **Low CHIR > No CHIR (339 transcripts*)** | | | |
| cell cycle | 85 | 2.4E-55 | 8.7 |
| cell division | 64 | 1.2E-46 | 10.8 |
| mitotic nuclear division | 57 | 5.9E-46 | 13.0 |
| chromosome segregation | 23 | 1.6E-20 | 16.3 |
| cellular response to DNA damage stimulus | 40 | 3.2E-19 | 6.0 |
| **No CHIR > Low CHIR (233 transcripts*)** | | | |
| cell adhesion | 8 | 1.8E-03 | 4.6 |
| cell migration | 7 | 2.4E-03 | 5.1 |
| cell cycle arrest | 5 | 3.1E-03 | 8.2 |
| negative regulation of cell proliferation | 9 | 4.8E-03 | 3.4 |
| regulation of cell growth | 4 | 4.8E-03 | 11.5 |

* These genes were identified with a different DE threshold.
