## Supplemental Figure 2 for "A β-catenin-driven switch in TCF/LEF transcription factor binding to DNA target sites promotes commitment of mammalian nephron progenitor cells"

Fig. S2

A

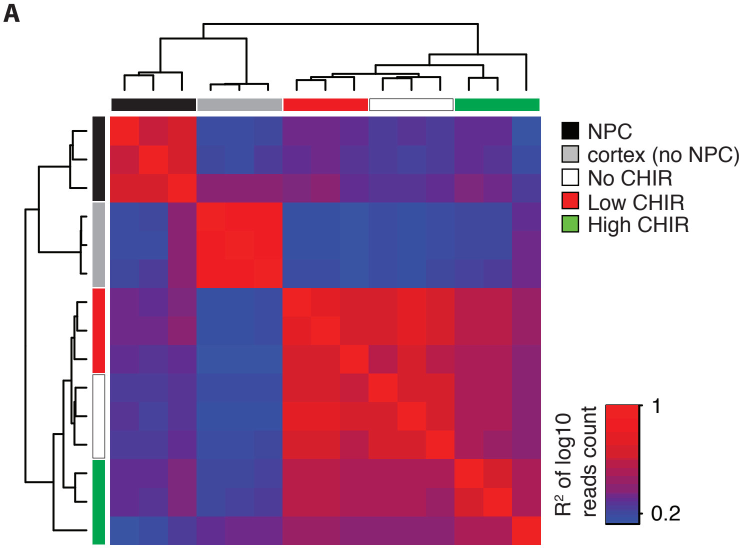


B

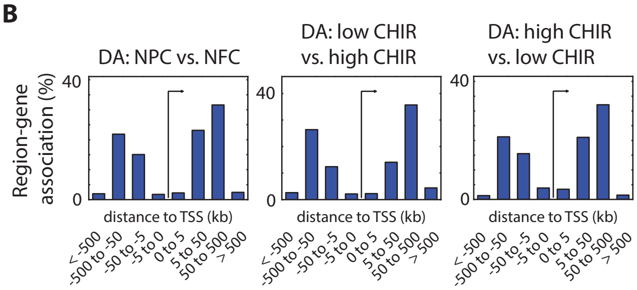


C

**NFC vs. NPC: 2482 regions**

| GO terms | FDR | f.c. | region # | gene # |
|---|---|---|---|---|
| cardiac chamber morphogenesis | 9E-09 | 2.1 | 90 | 42 |
| cardiac chamber development | 3E-08 | 2.0 | 97 | 47 |
| osteoblast differentiation | 4E-08 | 2.5 | 60 | 29 |
| outflow tract morphogenesis | 3E-07 | 2.3 | 63 | 23 |
| cardiac ventricle development | 3E-07 | 2.0 | 84 | 37 |

D

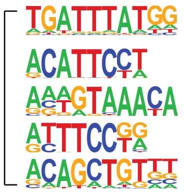


| motif logo | p-value | pred. TF |
|---|---|---|
| | 1e-81 | Hox |
| | 1e-45 | Tead |
| | 1e-40 | Fox |
| | 1e-36 | Etv |
| | 1e-35 | bHLH |
