## Supplementary figures and images for "A β-catenin-driven switch in TCF/LEF transcription factor binding to DNA target sites promotes commitment of mammalian nephron progenitor cells"

### Supplemental Figure 3

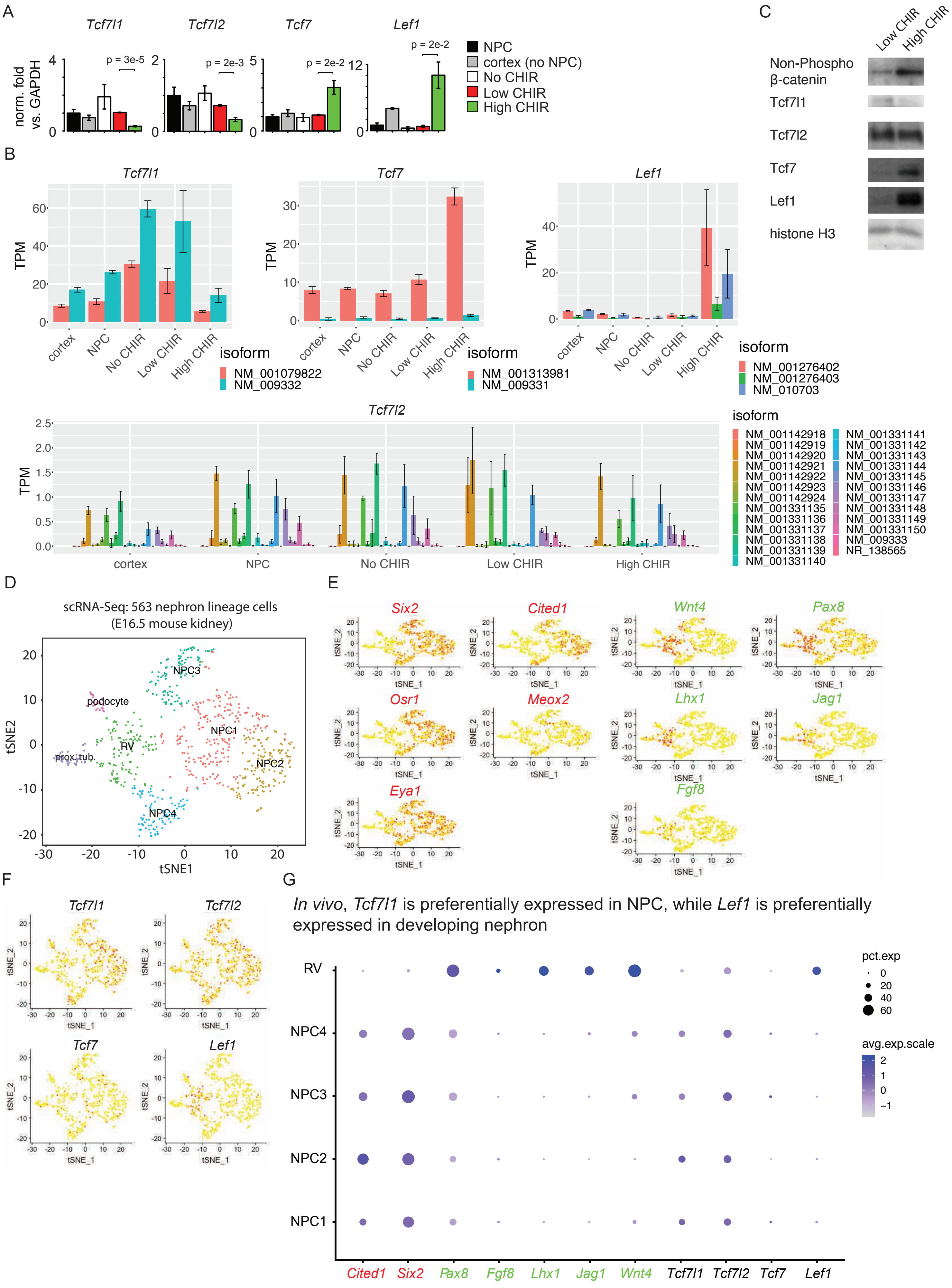

### Supplemental Figure 5

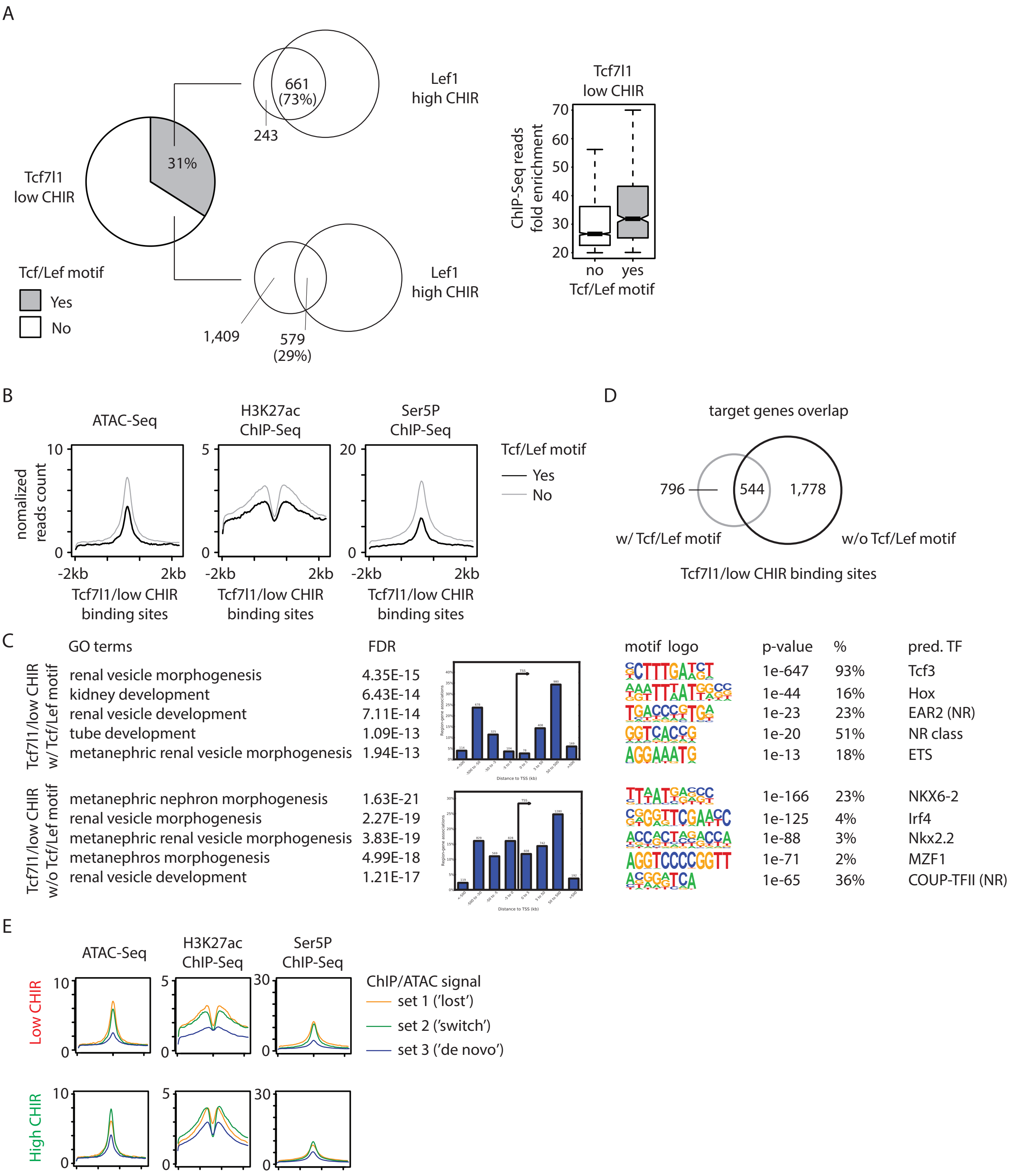

### Supplemental Figure 6

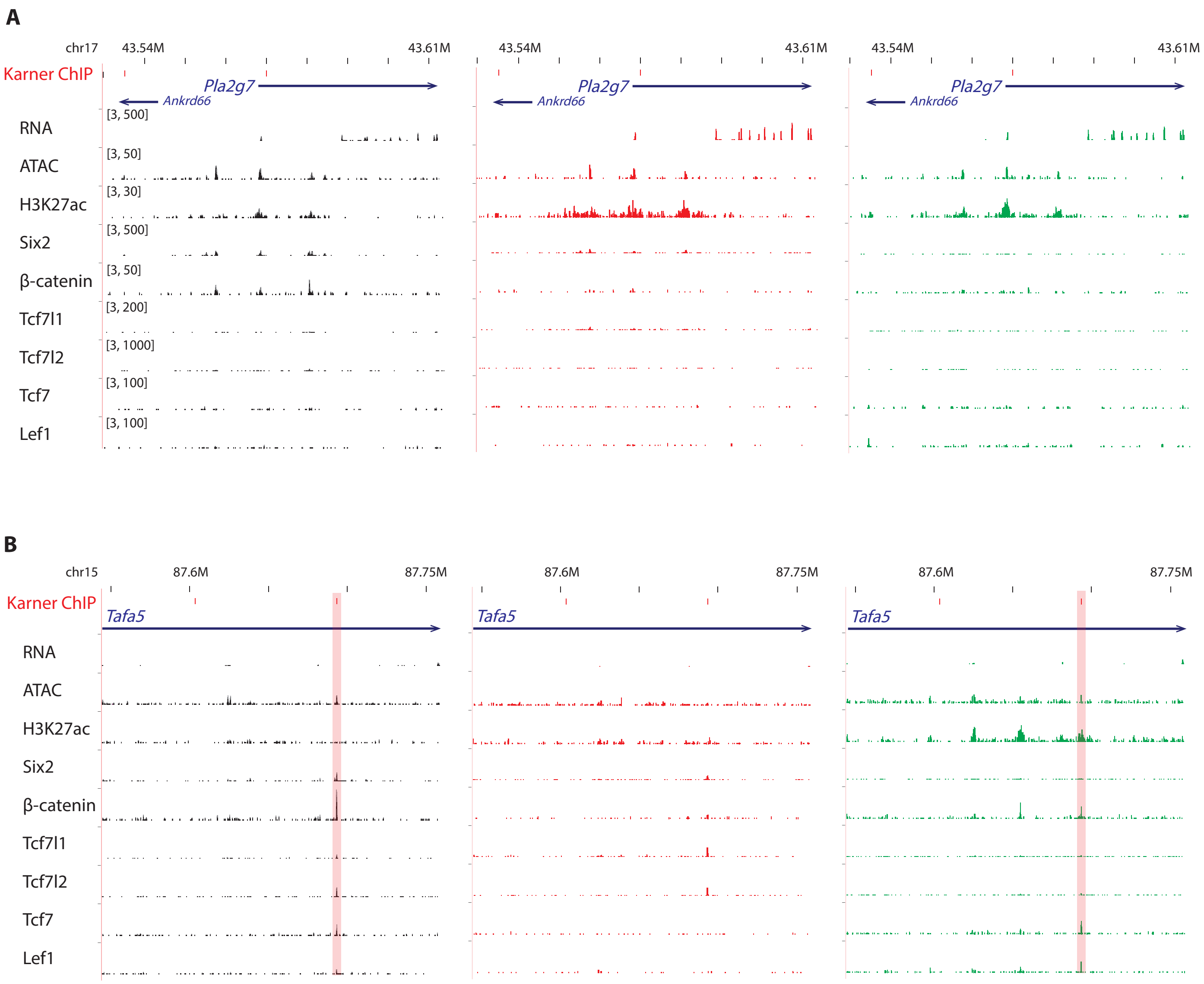

### Supplemental Figure 7

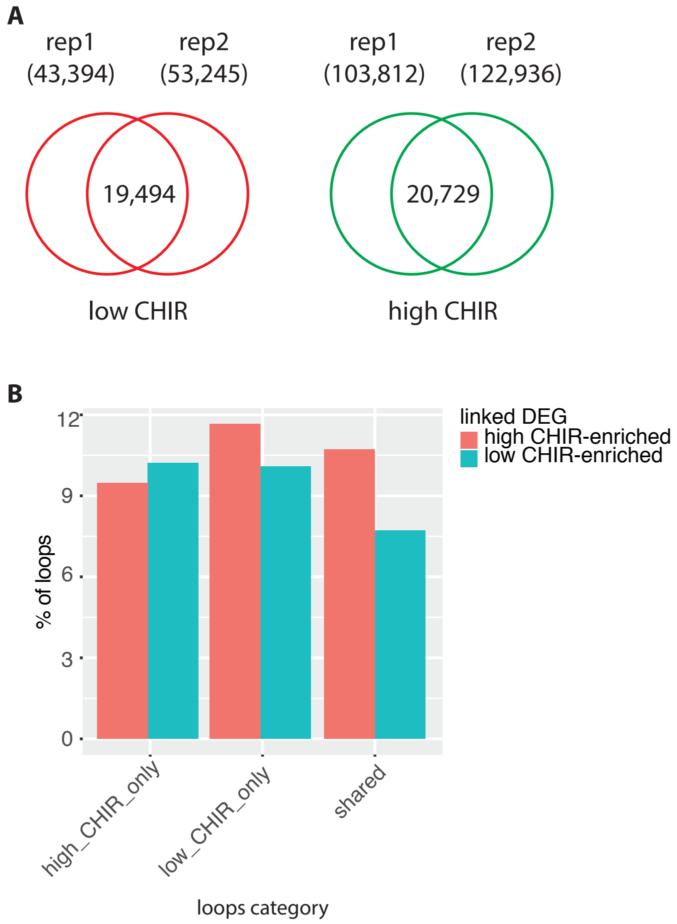
