## Supplemental Figure 4 for "A β-catenin-driven switch in TCF/LEF transcription factor binding to DNA target sites promotes commitment of mammalian nephron progenitor cells"

Fig. S4

**A**

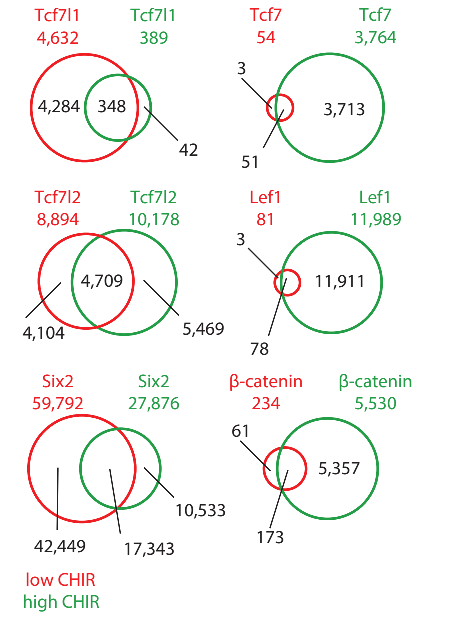


**B**

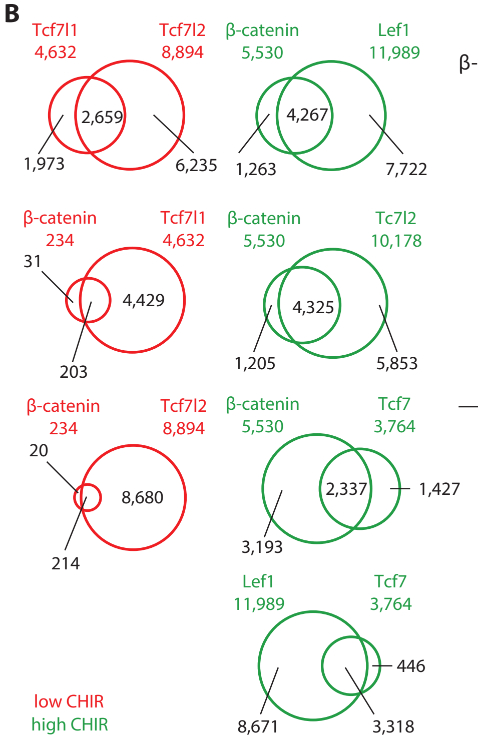


**C**

| | low CHIR | | | high CHIR | | |
|---|---|---|---|---|---|---|
| **Tcf7l1** | # sites = 4,632 | | | # sites = 389 | | |
| | Tcf/Lef | 1e-401 | 19.2/3.2 | NA | NA | NA |
| | Nkx/Hox | 1e-195 | 12.5/2.8 | | | |
| | Irf4* | 1e-132 | 2.5/0.1 | | | |
| | Coup-TFII | 1e-83 | 25.7/14.8 | | | |
| | Barx1/Sox | 1e-71 | 0.7/0.0 | | | |
| **Tcf7l2** | # sites = 8,894 | | | # sites = 10,178 | | |
| | Tcf/Lef | 1e-2224 | 39.2/4.5 | Tcf/Lef | 1e-3517 | 44.0/3.6 |
| | Nkx/Hox | 1e-346 | 22.4/8.5 | Nkx/Hox | 1e-486 | 19.7/5.9 |
| | Coup-TFII | 1e-199 | 26.3/14.1 | Runx | 1e-199 | 9.3/2.8 |
| | Six2 | 1e-112 | 8.7/3.5 | Coup-TFII | 1e-112 | 15.6/7.2 |
| | Wt1 | 1e-93 | 5.2/1.7 | Hox | 1e-93 | 4.8/1.5 |
| **Tcf7** | # sites = 55 | | | # sites = 3,764 | | |
| | Tcf/Lef | 1e-67 | | Tcf/Lef | 1e-1984 | 51.1/2.5 |
| | | | | Runx* | 1e-136 | 6.7/0.8 |
| | | | | Nkx/Hox | 1e-108 | 12.5/3.8 |
| | | | | Coup-TFII* | 1e-70 | 14.0/6.0 |
| **Lef1** | # sites = 81 | | | # sites = 11,989 | | |
| | Tcf/Lef | 1e-49 | | Tcf/Lef | 1e-4535 | 37.2/1.7 |
| | | | | Hox | 1e-383 | 16.6/5.8 |
| | | | | Runx | 1e-227 | 8.5/2.6 |
| | | | | AP-1 | 1e-75 | 4.9/2.1 |
| | | | | Pax8 | 1e-61 | 3.0/1.1 |
| | | | | Ctcf | 1e-51 | 1.8/0.5 |
| **β-catenin** | # sites = 234 | | | # sites = 5,530 | | |
| | Tcf/Lef | 1e-45 | | Tcf/Lef | 1e-1057 | 37.4/5.8 |
| | Hox | 1e-28 | | Nkx/Hox | 1e-469 | 26.4/6.5 |
| | | | | Runx | 1e-91 | 7.4/2.3 |
| | | | | Pax2 | 1e-90 | 25.3/14.9 |
| | | | | AP-1 | 1e-83 | 8.1/2.8 |
| **Six2** | # sites = 59,792 (top 10,000) | | | # sites = 27,876 (top 10,000) | | |
| | Six2 | 1e-4319 | 49.0/3.3 | Six2 | 1e-3724 | 34.1/1.3 |
| | Meis1 | 1e-447 | 66.0/43.5 | Six4 | 1e-429 | 15.0/3.9 |
| | Nkx/Hox | 1e-299 | 8.7/1.8 | GFY | 1e-335 | 3.5/0.2 |
| | Wt1 | 1e-134 | 7.5/2.6 | Nkx/Hox | 1e-284 | 9.6/2.4 |
| | Etv | 1e-111 | 19.6/11.7 | Tcf/Lef | 1e-140 | 7.2/2.4 |

**D**

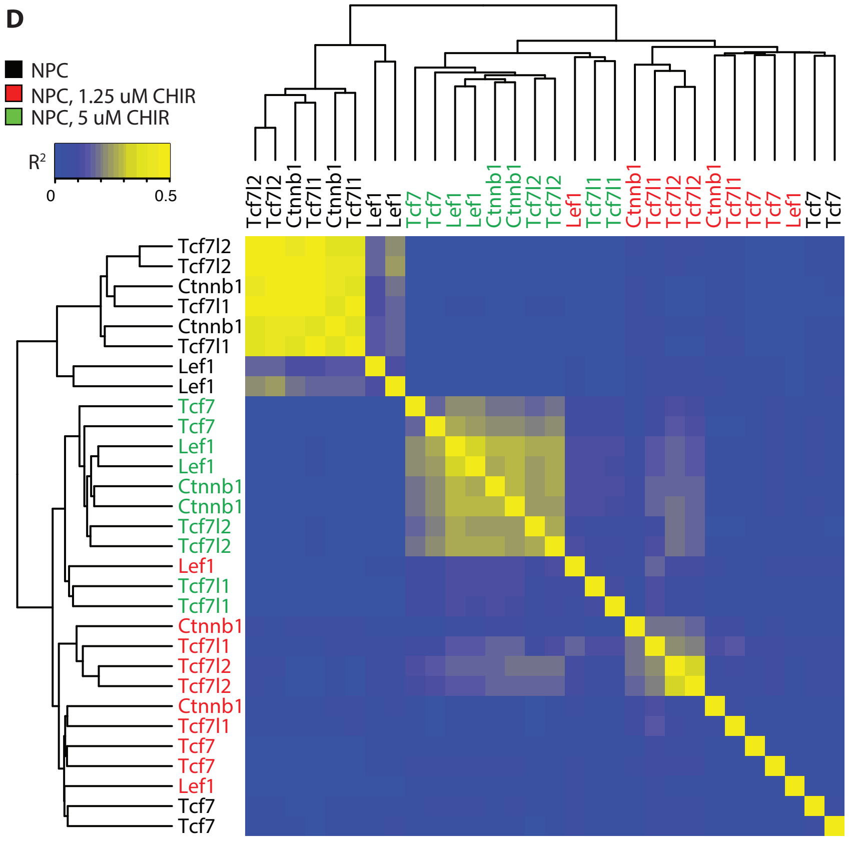
